# Paired CRISPR screens to map gene regulation in *cis* and *trans*

**DOI:** 10.1101/2024.11.27.625752

**Authors:** Xinhe Xue, Zoran Z. Gajic, Christina M. Caragine, Mateusz Legut, Conor Walker, James Y.S. Kim, Xiao Wang, Rachel E. Yan, Hans-Hermann Wessels, Congyi Lu, Neil Bapodra, Gamze Gürsoy, Neville E. Sanjana

## Abstract

Recent massively-parallel approaches to decipher gene regulatory circuits have focused on the discovery of either *cis*-regulatory elements (CREs) or *trans*-acting factors. Here, we develop a scalable approach that pairs *cis*- and *trans*-regulatory CRISPR screens to systematically dissect how the key immune checkpoint *PD-L1* is regulated. In human pancreatic ductal adenocarcinoma (PDAC) cells, we tile the *PD-L1* locus using ∼25,000 CRISPR perturbations in constitutive and IFNγ-stimulated conditions. We discover 67 enhancer- or repressor-like CREs and show that distal CREs tend to contact the promoter of *PD-L1* and related genes. Next, we measure how loss of all ∼2,000 transcription factors (TFs) in the human genome impacts PD-L1 expression and, using this, we link specific TFs to individual CREs and reveal novel PD-L1 regulatory circuits. For one of these regulatory circuits, we confirm the binding of predicted *trans*-factors (SRF and BPTF) using CUT&RUN and show that loss of either the CRE or TFs potentiates the anti-cancer activity of primary T cells engineered with a chimeric antigen receptor. Finally, we show that expression of these TFs correlates with *PD-L1* expression *in vivo* in primary PDAC tumors and that somatic mutations in TFs can alter response and overall survival in immune checkpoint blockade-treated patients. Taken together, our approach establishes a generalizable toolkit for decoding the regulatory landscape of any gene or locus in the human genome, yielding insights into gene regulation and clinical impact.

---

Alterations in regulators of gene expression underlie diverse human diseases^1^, including cancer^2^. *Cis*- regulatory elements (CREs), such as promoters, enhancers, silencers and insulators, govern gene expression but do not themselves directly code for proteins^3^. Instead, they are activated by the binding of *trans*- regulatory machinery including transcription factors (TFs) and chromatin remodelers^2^. These regulatory complexes interact with target gene promoters via 3D looping to orchestrate gene expression with remarkable specificity^4^. Recent high-throughput methods for protein-coding gene knock-out and overexpression have sped the identification of these *trans*-regulators for many disease associated genes^5,6^. In a similar fashion, CRISPR-mediated mutagenesis of noncoding genomic loci have uncovered key CREs within individual genomic loci^7–9^ — including the discovery of a noncoding regulator of fetal hemoglobin where mutagenesis can cure sickle-cell anemia and beta-thalessemia^10^. Despite these remarkable advances, there is a lack of systematic integration of *cis*- and *trans*-regulator discovery to comprehensively map the regulatory landscape of gene expression.

Across multiple different cancers, the expression of certain tumor ligands can predict clinical outcomes to certain immunotherapies such as immune checkpoint blockade (ICB) therapies^11–13^. For example, the expression of the inhibitory immune ligand PD-L1, which binds the PD-1 receptor on the surface of T cells and other immune cells^14,15^, predicts response to ICB therapies and is an approved companion diagnostic^13,16,17^. Engagement between PD-L1 and PD-1 impairs T cell activation and allows tumor cells to escape immune surveillance. Hence, understanding of the complete regulatory landscape — including both *cis*- and *trans*-regulators — for such ligands is of paramount importance. To date, there have been no efforts to systematically map noncoding CREs that control *PD-L1* expression in tumors.

Here, using a saturation mutagenesis CRISPR screen tiling ∼1 megabase surrounding the *PD-L1* locus in PDAC cells, we discover 67 enhancer- and repressor-like candidate CREs (cCREs) governing PD-L1 constitutive and IFNγ-stimulated expression with context-specific activity. For a subset of these cCREs, we quantify how their loss impacts PD-L1 expression and identify 3D interactions with target gene promoters, suggesting that transcription factors (TFs) bound to these distal sites act directly to modulate PD-L1 expression. To identify likely TFs at the PD-L1 CREs, we perform a knock-out screen targeting all TFs in the human genome. By integrating six CRE- and TF-focused CRISPR screens, we pair 13 validated CREs of PD-L1 to one or more PD-L1 *trans*-regulators. For one of these regulatory modules, we demonstrate direct binding of two *trans*-regulators at the CRE and show that loss of either the *cis*-element or *trans*- regulator alters the chimeric antigen receptor (CAR)-T cytotoxicity and tumor immune evasion. Lastly, we find that expression of some PD-L1 *trans*-regulators — including almost all activator regulators — correlate with PD-L1 expression across 178 PDAC tumors. One example is the PD-L1 repressor *FOXA3*:

It is recurrently mutated in PDAC patients and mutations in *FOXA3* impact the responsiveness and overall survival of ICB-treated patients. These paired CRISPR screens and complementary analyses provide a deep dissection of the regulatory networks modulating PD-L1 expression.

## Saturation mutagenesis maps *cis*-regulatory elements of *PD-L1*

To identify CREs at the *PD-L1* locus, we designed a library of 25,537 CRISPR-Cas9 guide RNAs (gRNAs) to tile the 768 kb topologically-associated domain (TAD) surrounding *PD-L1* (**Fig. 1a**, **Supplementary Fig. 1a, b, Supplementary Tables 1-3**). This library achieves near-saturation mutagenesis over the entire locus; the median distance between cut sites is 12 bp and repair after Cas9 cutting results in, on average, ∼10 bp indels^8^ (**Supplementary Fig. 1c**). We lentivirally transduced BxPC-3 pancreatic adenocarcinoma cells at a low multiplicity of infection so that each cell would receive one perturbation. We maintained the cells at a high representation (∼1000 cells per perturbation) and waited 14 days after lentiviral transduction for complete genome modification. Next, we labeled cells with a PD-L1 antibody and sorted cells by their PD-L1 expression, isolating the top and bottom 20% populations (**Supplementary Fig. 1d**). We found that gRNAs targeting PD-L1 coding exons — which result in PD-L1 loss — were highly enriched in PD-L1^low^ sorted cells and highly depleted in PD-L1^high^ cells (**Supplementary Fig. 1e-i**), serving as a strong positive control for the phenotypic selection.

**Figure 1.**
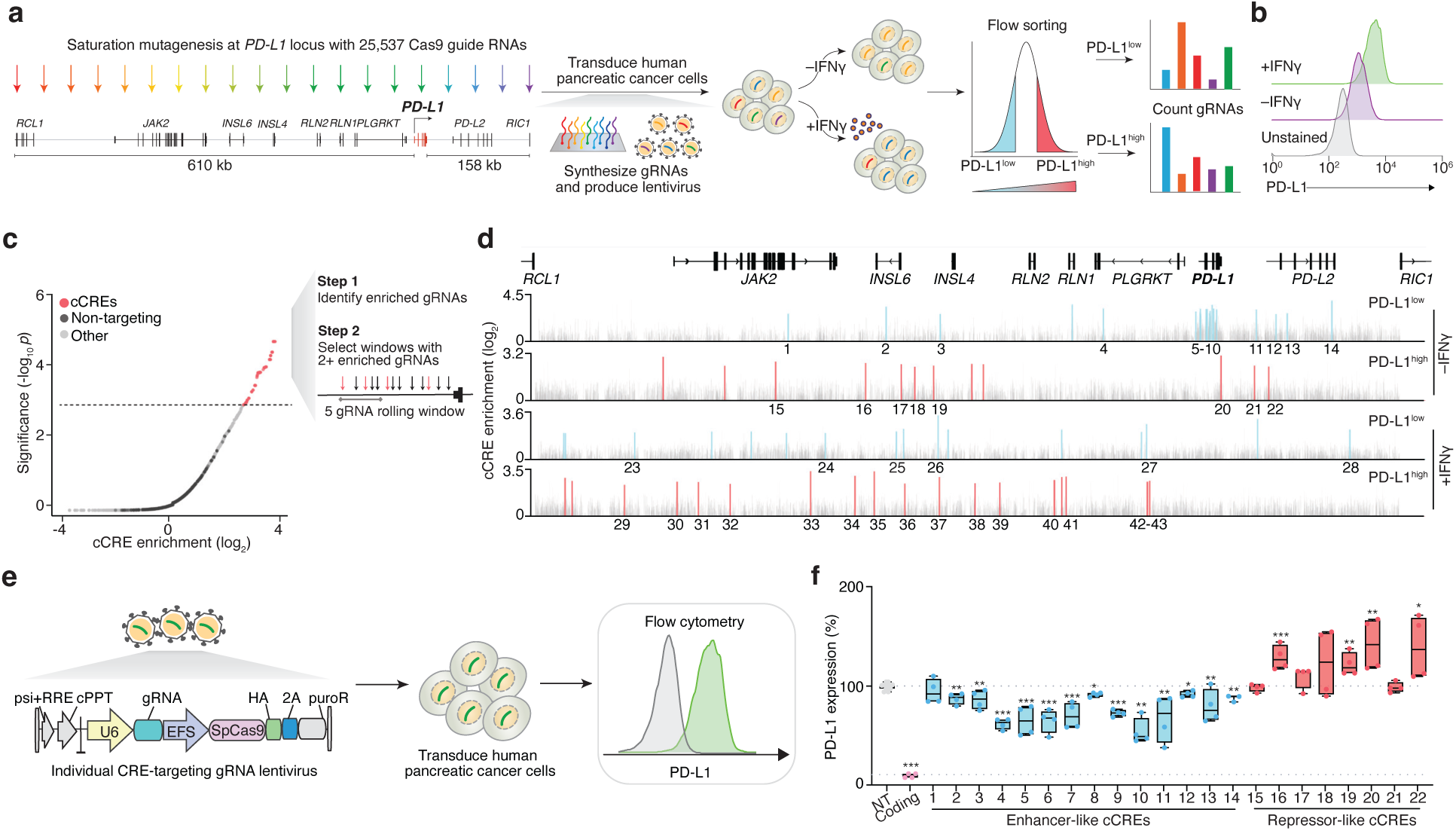
A saturation mutagenesis CRISPR screen to map *cis*-regulatory elements of *PD-L1*. **a,** A pooled saturation mutagenesis screen of a ∼800 kb topologically-associated domain containing the *PD-L1* locus using ∼25,000 CRISPR guide RNAs (gRNAs). **b,** *PD-L1* expression in untreated and interferon (IFNγ)-treated human BxPC-3 pancreatic ductal adenocarcinoma cells. **c,** Window-based consistency metric to identify candidate *cis*-regulatory elements (cCREs) at the *PD-L1* locus. **d,** Enhancer-like cCREs (enriched in PD-L1^low^ in *blue*) and repressor-like cCREs (enriched in PD-L1^high^ in *red*) for constitutive or IFNγ-stimulated PD-L1 expression identified using the approach in panel *c* (*n* = 67 cCREs). The cCRE enrichment is the fold-change of CRE-targeting gRNA abundance in sorted cell populations compared to presort cell populations. The numbered cCREs were selected for arrayed validation. Windows without cCREs are indicated in *grey*. **e,** Arrayed validation of individual cCREs to measure change in PD-L1 expression. **f,** PD-L1 expression after transduction with individual non-targeting (NT), *PD-L1* coding exon-targeting (Coding), enhancer-like and repressor-like cCRE-targeting gRNAs (*n* = 2 - 4 biological replicates of 2-3 gRNAs each). cCRE-6 and cCRE-7 are resided at *PD-L1* promoter. Expression values are normalized to NT group and *p*-values denote Bonferroni-adjusted Mann-Whitney *U* test to NT gRNA-transduced cells: \**p* < 0.05, \*\**p* < 0.01, \*\*\**p* < 0.001.

PD-L1 expression is regulated differently under constitutive and IFNγ-stimulated conditions in tumor cells^18,19^ and we observed a ∼3-fold increase in PD-L1 upon IFNγ treatment in BxPC-3 pancreatic adenocarcinoma cells (**Fig. 1b**, **Supplementary Fig. 1j**). To map differences in PD-L1 regulation after IFNγ stimulation, we treated library-transduced cells with IFNγ for 2 days and performed the same sorting (**Supplementary Fig. 1k**). After IFNγ stimulation, we found that gRNAs that target *JAK2* exons were enriched in PD-L1^low^ cells since JAK2 is required for IFNγ-driven PD-L1 expression^20,21^ (**Supplementary Fig. 1l-o**). As expected, we did not see any enrichment of gRNAs targeting *JAK2* exons under constitutive conditions (**Supplementary Fig. 1f-i**). Although the CRISPR library included exon-targeting gRNAs for all 9 genes in the locus, only *JAK2*, *PD-L1,* and *PD-L2* gRNAs were enriched in IFNγ-treated PD-L1^low^ cells, indicating that *JAK2* and *PD-L2* can contribute to *PD-L1* regulation in IFNγ-stimulated conditions. As negative controls, we also included 1,000 gRNAs designed not to target anywhere in the human genome and, on average, these non-targeting gRNAs were slightly depleted in each flow-sorted population (**Supplementary Fig. 1g, m**). Taken together, these positive and negative control gRNAs validate our approach to detecting changes in PD-L1 expression in both constitutive and IFNγ-stimulated conditions, enabling us to look for noncoding regulators of expression.

We identified PD-L1 CREs based on consistent enrichment of gRNAs that target nearby regions of the genome^22^ (**Fig. 1c**, **Supplementary Fig. 2a, b, Supplementary Table 4**). In total, we identified 67 candidate CREs (cCREs) governing PD-L1 expression over the ∼800 kb region targeted (**Fig. 1d**, **Supplementary Fig. 2c, d, Supplementary Table 5**). We found that cCREs detected in PD-L1^low^ and PD- L1^high^ populations were distinct from each other — there was no overlap in cCREs identified in PD-L1^low^ and PD-L1^high^ populations, highlighting a clear separation between enhancer- and repressor-like cCREs. We also observed differences in cCRE locations: In constitutive conditions, the average repressor-like cCRE is located more distally from the *PD-L1* transcription start site (TSS) than the average enhancer-like cCRE (∼254 kb for repressor-like versus ∼73 kb for enhancer-like cCREs) (**Supplementary Fig. 2e**). Similarly, no cCREs were shared between constitutive and IFNγ-stimulated conditions for either enhancer- or repressor-like cCREs.

Promoter regions and introns tend to harbor gene regulatory elements^7–10^ and, accordingly, we found many enhancer-like cCREs in cells without IFNγ treatment near the *PD-L1* promoter and initial introns (see cCREs 5 – 10, **Fig. 1d**). However, upon IFNγ treatment, we observed a striking loss of PD-L1 regulation by these proximal elements. In IFNγ-treated cells, we identify several PD-L1 cCREs proximal to *JAK2* (e.g. cCRE 23), which is expected since increased JAK2 drives increases in PD-L1 through IFNγ-induced JAK-STAT activation^21,23^.

Recently, two studies have identified a super-enhancer^24^ and a 3’ UTR^25^ variant that impact PD-L1 expression. Xu et al. identified an 8 kb super-enhancer located between PD-L1 and PD-L2^24^. In our screens, this enhancer-like cCRE (cCRE 11) also drives PD-L1 expression. However, with our high-resolution tiling, we were able to refine this cCRE to a smaller region: We identify a ∼100 bp core region, yielding a 80-fold increase in resolution and highlighting an advantage of dense mutagenesis. Another study by Kataoka et al. identified a truncated 3’ UTR transcript variant that drives elevated PD-L1 levels^25^. We found that mutagenesis at the DNA-level in the same location within the 3’ UTR also leads to PD-L1 overexpression (cCRE 20). Here, our tiling approach detects a smaller region (∼50 bp) within the larger 4 kb deletion used by Kataoka et al. The ability to detect (and refine) established noncoding regulators bolsters our confidence in the detected cCREs from the saturation mutagenesis screens.

To confirm that mutagenesis of these cCREs modulates PD-L1 expression, we individually perturbed cCREs with two gRNAs each and measured PD-L1 surface protein expression after mutagenesis (**Fig. 1e**). In the constitutive state, we found that mutagenesis of 13 out of 14 enhancer-like cCREs tested significantly reduced PD-L1 compared to non-targeting gRNA-transduced cells (**Fig. 1f**, **Supplementary Fig. 2f**). Mutagenesis of proximal CREs (such as those in the introns of *PD-L1*) had a larger impact on PD-L1 expression than more distal CREs (**Supplementary Fig. 2g**), in line with prior studies^8^. We also identified 4 repressor-like CREs that significantly increased PD-L1 expression after CRE mutagenesis (**Fig. 1f**). In IFNγ-treated cells, we found that all 6 of the enhancer-like cCREs that we assayed led to significant PD-L1 reduction and that 3 repressor-like CREs yielded an increase in PD-L1 after mutagenesis (**Supplementary Fig. 2h**).

## Most PD-L1 CREs contact the *PD-L1* or *JAK2* promoter

Given that IFNγ stimulation results in different PD-L1 CREs, we wondered whether changes in 3D genome folding may be responsible for bringing different sets of distal regulatory elements into close proximity with the promoter of *PD-L1* and related gene *JAK2*^21,23^. To measure this in constitutive and IFNγ-stimulated conditions, we performed a modified version of H3K27ac-anchored contact mapping that uses a fixed spike-in of mouse nuclei for absolute quantification of chromatin architecture (AQuA-HiChIP) (**Supplementary Fig. 3a**). We first verified that there was a higher percentage of *cis*-interactions (within the same chromosome) than *trans*-interactions (different chromosomes) as observed in prior AQuA-HiChIP studies^26^ (**Supplementary Fig. 3b**).

Overall, we found a decrease in 3D contacts after IFNγ treatment, within the *PD-L1* TAD and across the genome (**Fig. 2a-b**, **Supplementary Fig. 3c-h, Supplementary Table 6**). To investigate promoter-enhancer interactions, we examined H3K27ac HiChIP loops and found that a third of the HiChIP loops in the *PD-L1* TAD contact a gene promoter in both constitutive and IFNγ-stimulated conditions (**Supplementary Fig. 3i)**. Nearly half of the promoter loops in the *PD-L1* TAD contact *PD-L1* or the known PD-L1 regulator *JAK2*. In line with the global reduction in loop strength after IFNγ treatment, we also observed a reduced strength of promoter-anchored loops in the *PD-L1* TAD upon IFNγ treatment (**Fig. 2c**, **Supplementary Fig. 3j**).

**Figure 2.**
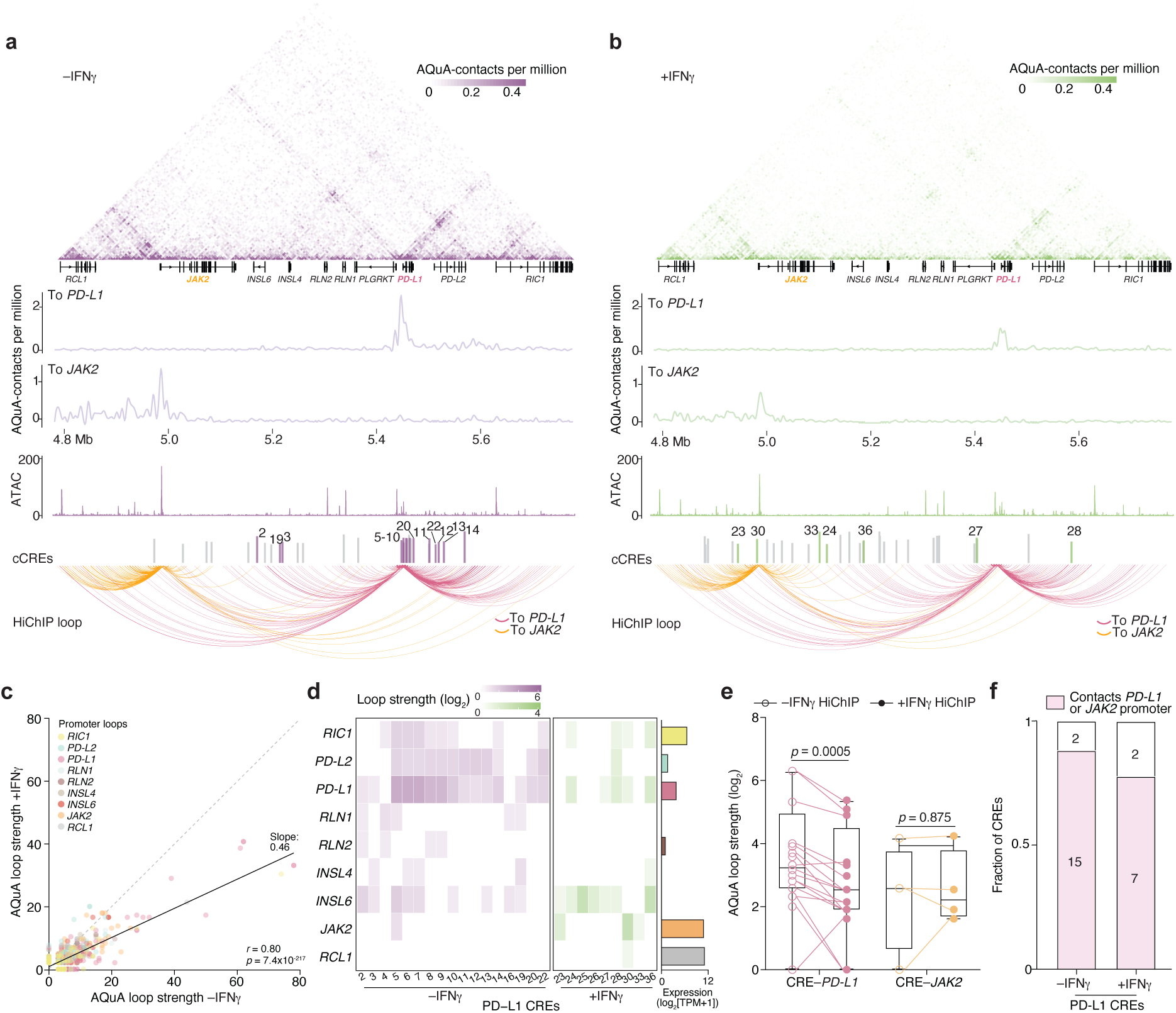
H3K27ac HiChIP reveals physical contacts between PD-L1 CREs and gene promoters within the *PD-L1* TAD. **a, b,** Chromatin architecture in untreated (*a*) and IFNγ-treated (*b*) BxPC-3 cells. *Upper plot*: AQuA- normalized valid HiChIP contact map at 5-kb resolution. *Bottom plot*: HiChIP loops to the promoter of *PD- L1* (*pink*) and *JAK2* (*orange*). The thickness of the HiChIP loops indicate the AQuA-normalized loop strength. Validated CREs that contact the promoter of *PD-L1* and *JAK2* are numbered and colored. **c,** Spearman correlation of promoter-anchored HiChIP loops in the *PD-L1* TAD between constitutive and IFNγ-stimulated contexts. **d,** Contact strength between PD-L1 CREs and indicated genes in the *PD-L1* TAD. For each gene, expression in BxPC-3 cells is shown (*right*). **e,** HiChIP loop strength for loops between PD-L1 CREs and the promoter of *PD-L1* or *JAK2* in constitutive and IFNγ-stimulated contexts. Significance was determined by a Wilcoxon signed rank test. **f,** Overview of PD-L1 CREs with physical contacts to the promoters of *PD-L1* or *JAK2*.

Focusing on the PD-L1 CRISPR screen CREs, we observed most of the PD-L1 CREs contacted more than one promoter — with *PD-L1* and *PD-L2* being the most common pleiotropy (**Fig. 2d**). For example, a previously reported super-enhancer of *PD-L1* and *PD-L2*^24^ (CRE11) contacts both promoters with similar contact strength. We found that the change in PD-L1 protein expression upon CRISPR mutagenesis of the PD-L1 screen CREs correlated well with the strengths of the CRE*-PD-L1* promoter loops — but not with the strengths of CRE-*PD-L2* promoter loops (**Supplementary Fig. 3k, l**).

Separating the CRISPR screen CREs by condition (constitutive and IFNγ-stimulated), we found that the majority of CREs in the constitutive condition contact the promoter of *PD-L1*, whereas, in the IFNγ stimulation condition, CREs tended to contact the promoter of either *PD-L1* or *JAK2* (**Fig. 2d**, **Supplementary Fig. 3m**). Notably, CRE-promoter looping decreased upon IFNγ stimulation to *PD-L1* but not to *JAK2* (**Fig. 2e**), which modulates PD-L1 expression via IFNγ-induced JAK-STAT signaling^21,23^. For example, CRE33 only contacts *JAK2* upon IFNγ stimulation and likely regulates PD-L1 expression indirectly via JAK2 modulation. Overall, we found that ∼90% of validated PD-L1 CREs directly contact the promoter of *PD-L1* or *JAK2* (**Fig. 2f**). This suggests that recruitment of *trans*-factors (e.g. TFs) to these CREs likely places them in close 3D proximity to promoters for effective gene regulation and that the positions of CREs change dynamically upon IFNγ stimulation.

## A TFome-wide CRISPR screen identifies established and novel *trans*-regulators for PD-L1 expression

We next sought to identify which TFs are important for constitutive and IFNγ-stimulated PD-L1 expression in PDAC cells. To comprehensively map TFs that influence PD-L1 expression, we performed TFome-wide CRISPR knock-out screens targeting 1,891 human TFs with 10 gRNAs per TF^27^ (**Fig. 3a**, **Supplementary Fig. 4a, Supplementary Table 7**). First, we lentivirally transduced BxPC-3 PDAC cells with TF-targeting gRNAs in replicates and measured constitutive and IFNγ-stimulated PD-L1 expression using flow cytometry (**Supplementary Fig. 4b, c**). Then, we collected cells with increased or reduced PD-L1 expression, enabling the identification of both activators and repressors of PD-L1, and did so in both untreated and IFNγ-treated cells.

**Figure 3.**
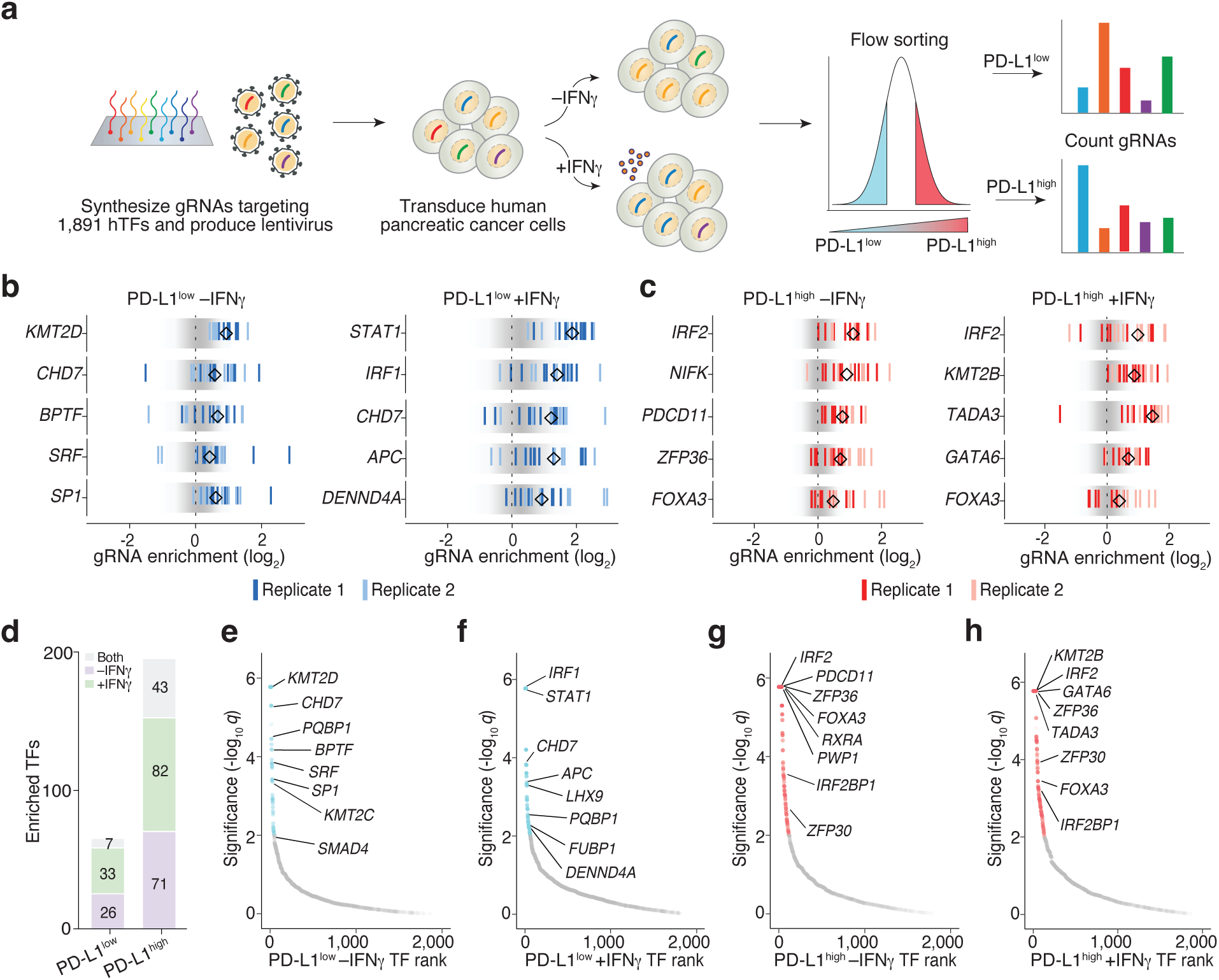
A TFome-wide CRISPR loss-of-function screen identifies known and novel *trans*-regulators of PD-L1 expression. **a,** A pooled CRISPR screen for PD-L1 protein expression that targets all human transcription factors (TFs) with 10 guide RNAs (gRNAs) each in human BxPC-3 pancreatic ductal adenocarcinoma cells. **b,** Normalized enrichment of gRNAs for the indicated TFs in untreated (*left*) and IFNγ-treated (*right*) PD- L1^low^ flow-sorted cells (*n* = 10 guide RNAs per gene in each of 2 biological replicates). The diamond denotes the median gRNA enrichment across biological replicates and the distribution of the 1,000 negative control (non-targeting) gRNAs is shown in *grey*. The gRNA enrichment is the fold-change of gRNA abundance in PD-L1^low^ cell populations compared to presort cell populations. **c,** Normalized enrichment of gRNAs for the indicated TFs in untreated (*left*) and IFNγ-treated (*right*) PD- L1^high^ flow-sorted cells (*n* = 10 guide RNAs per gene in each of 2 biological replicates). The diamond denotes the median gRNA enrichment across biological replicates and the distribution of the 1,000 negative control (non-targeting) gRNAs is shown in *grey*. The gRNA enrichment is the fold-change of gRNA abundance in PD-L1^high^ cell populations compared to presort cell populations. **d,** Activator and repressor *trans*-regulators for *PD-L1* expression that are shared across constitutive and IFNγ-stimulated conditions or unique to each condition. **e-h,** Robust rank aggregation (RRA) of TFs that modulate PD-L1 expression in untreated (*e*) and IFNγ- treated (*f*) PD-L1^low^ flow-sorted cells, and in untreated (*g*) and IFNγ-treated (*h*) PD-L1^high^ flow-sorted cells. FDR-adjusted RRA *q*-values are derived from consistent enrichment of gRNAs targeting the same TF (*n* = 10 gRNAs per TF and 2 biological replicates of each pooled screen).

We identified several TFs in each sorted population with consistent enrichment between different gRNAs targeting the same TF (**Fig. 3b, c**, **Supplementary Fig. 4d)** and further confirmed these genes are robustly expressed in BxPC-3 cells (**Supplementary Fig. 4e, Supplementary Table 8**). Next, we ranked each of the 1,891 TFs in the four sorted populations. As a result, we identified 66 activator and 196 repressor *trans*- regulators for PD-L1 expression which are significantly enriched in the sorted populations (FDR-adjusted RRA *q* ≤ 10^-2^) and the majority of which are specific to each sorted population (**Fig. 3d-h**).

Reassuringly, several of the top-ranked TFs in each population have previously been reported to play a role in PD-L1 expression. For example, SP1, which binds the promoter of *PD-L1* and upregulates its expression in gastric cancer^28^, was ranked highly in non-IFNγ-treated PDAC cells. KMT2C, another top-ranked activator TF, has previously been suggested as a biomarker for anti-PD-1 checkpoint blockade therapy in melanoma^29^. In IFNγ-treated cells, IRF1 and STAT1 were the top activator regulators of PD-L1, which is expected given the role of the JAK-STAT pathway in driving PD-L1 expression^21,30^.

We next validated several highly-ranked TFs and measured how their loss impacts PD-L1 expression in different contexts (**Supplementary Fig. 4f, g).** For example, IRF1 showed IFNγ-dependent regulation of PD-L1 expression with no effect on PD-L1 expression in the constitutive context. In contrast to IRF1 and STAT1, loss of IRF2 was previously shown to increase PD-L1 in multiple tumors and drive immune evasion^30,31^. In agreement, we identified IRF2 as a top-ranked repressor TF in both untreated and IFNγ- treated PDAC cells (**Supplementary Fig. 4f, g)**. Furthermore, we found that IRF2BP1, which interacts with IRF2 as an IRF2-dependent transcriptional repressor^31^, was also a strong repressor of PD-L1 expression.

In these TFome-wide pooled screens, we also included several gRNAs targeting PD-L1 CREs (i.e. noncoding regions) that we had identified from the saturation mutagenesis screen. We found that mutagenesis of several CREs (including CRE5, CRE 11, and CRE 24) resulted in a similar degree of PD- L1 modulation as the *trans*-regulators (**Supplementary Fig. 4h, i**). This suggests that loss of function in either the TF regulators or the targeted CREs leads to comparable changes in PD-L1 expression and that both elements (TF and CRE) likely function together in an integrated module.

## Integration of *cis*- and *trans*-CRISPR screens to identify CRE-TF regulatory modules

To systematically identify CRE-TF regulatory modules, we sought to identify TF binding sites within individual CREs where TFs might act to modulate *PD-L1* transcription^32^ using JASPAR^33^ and HOCOMOCO^34^ motif databases (**Supplementary Fig. 5a, Supplementary Table 9**). After doing so, we examined both the likelihood of a TF motif occurring within each CRE and the enrichment of the TFs in PD-L1 regulation from the TFome-wide CRISPR screens (**Fig. 4a**): We hypothesized that TFs with a binding site within a CRE and highly-ranked in the TFome-wide screen may modulate PD-L1 expression via the corresponding CREs from the saturation mutagenesis CRISPR screen.

**Figure 4.**
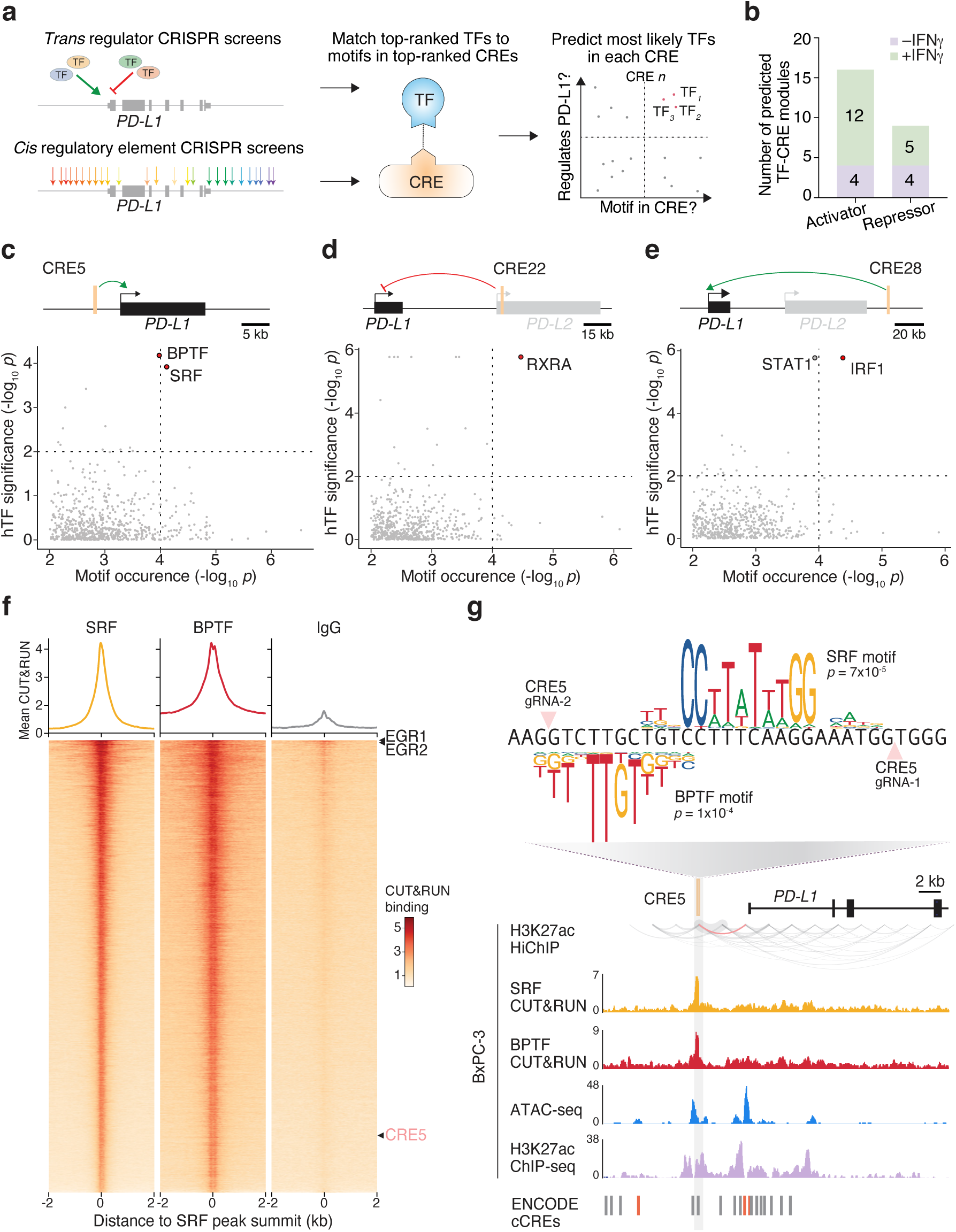
A subset of PD-L1 *trans*-regulators act on specific CREs for enhancing or repressing PD- L1 expression. **a,** Integration of *cis*- and *trans*-regulatory CRISPR screens to decode the regulatory landscape of PD-L1 expression. **b,** Predicted TF-CRE regulatory modules governing constitutive and IFNγ-stimulated PD-L1 expression. **c-e,** TF enrichment rank and motif identification for all TFs at: the enhancer-like CRE5 in the constitutive context (*c*), the repressor-like CRE22 in the constitutive context (*d*), and the enhancer-like CRE28 in the IFNγ-stimulated context (*e*). The *x*-axis shows the significance of TFs in PD-L1 regulation (*trans*- regulatory CRISPR screen). The *y*-axis indicates TF binding sites enriched within the indicated CRE (*cis*- regulatory CRISPR screen). The top panel shows the genomic locus for each CRE. **f,** CUT&RUN binding of SRF, BPTF and negative control (IgG) at SRF genome-wide binding sites, centered on SRF peak summit within a 4 kb window (*n* = 3 biological replicates). The mean CUT&RUN binding over all sites shown is plotted above. **g,** Genomic characterization of the CRE5 locus. *Upper plot*: The cut sites of two gRNAs targeting CRE5 and predicted motifs of SRF (JASPAR MA0083.2) and BPTF (HOCOMOCO H11MO.0.D) are indicated. *Bottom plot:* H3K27ac HiChIP contact map at 2-kb resolution (direct contacts between CRE5 and the promoter of *PD-L1* in *pink*), CUT&RUN binding of SRF and BPTF, ATAC-seq, H3K27ac ChIP and ENCODE CREs (Panc-1 pancreatic ductal adenocarcinoma in *red*).

By performing this integrative analysis at each PD-L1 CRE, we linked 13 validated CREs to at least one PD-L1 *trans*-regulator, yielding 25 putative TF-CRE regulatory modules involved in PD-L1 expression (**Fig. 4b**, **Supplementary Fig. 5b, Supplementary Table 10**). For example, at enhancer-like CRE5, we found the motifs of serum-response factor (SRF) and bromodomain PHD finger transcription factor (BPTF), which were top-ranked PD-L1 activator regulators in non-IFNγ-treated PDAC cells (ranked 8 and 9 out of 1,891 TFs) (**Fig. 4c**). We found similar interactions for repressor modulators of PD-L1: At repressor-like CRE22, we identified a binding motif for retinoid X receptor alpha (RXRA), which was a top-ranked PD- L1 repressor modulator (rank 26) (**Fig. 4d**). JAK2, STAT1 and IRF1 are canonical members of IFNγ- induced JAK-STAT signaling pathways^21^ and, at enhancer-like CRE28, we found an enrichment of the motifs for STAT1 and IRF1, which scored highly as activator regulators in IFNγ-stimulated cells (ranks 2 and 3 out of 1,891 TFs, respectively) (**Fig 4e**).

Although motif identification is suggestive of TF binding, we sought to examine if we could further validate a CRE-TF module by direct observation of TF binding at the CRE. To do this, we selected one enhancer-like CRE (CRE5) and performed CUT&RUN to identify binding sites genome-wide of SRF and BPTF, which were predicted by our integrative analysis to bind CRE5. We found that SRF colocalizes with BPTF throughout the genome, including at CRE5. Overall, 55% of SRF binding sites include BPTF within 200 nt — including at previously established SRF-BPTF binding sites at *EGR1* and *EGR2* loci^35^ (**Fig. 4f**, **Supplementary Table 11**).

Based on this data, we wondered whether the resulting indels created by CRE5-targeting gRNAs may disrupt SRF and BPTF binding sites. Indeed, we found that the top two enriched gRNAs for CRE5 were spaced close to the center of confirmed TF binding sites (7 nt and 21 nt from the SRF motif center and 10 nt and 18 nt from the BPTF motif center) (**Fig. 4g**). We found a similar pattern across many other CREs: For the 25 putative TF-CRE regulatory modules, we found that the CRE-targeting gRNAs whose cut sites were closer to the predicted TF binding sites exhibited greater changes in PD-L1 expression (**Supplementary Fig. 5c**). Using H3K27ac HiChIP, we also found looping between these CRE5 sites and the PD-L1 promoter, further reinforcing the mechanism by which recruitment of *trans*-regulators to CRE5 promotes *PD-L1* transcription (**Fig. 4g**). Besides CRE5, the majority of the PD-L1 regulatory modules (TF-CRE pairings) harbor CREs with HiChIP loops (11 out of 13 CREs) (**Supplementary Fig. 5b, d**). Thus, by combining TF motif enrichment analysis with the TFome-wide CRISPR screen, we can predict specific TF-CRE interactions that drive PD-L1 regulation.

## *Cis*- and *trans*-regulatory elements modulate tumor immune evasion

Given the prominent role of PD-L1 in T cell inhibition, we wondered whether changes in PD-L1 due to CRE mutagenesis or TF loss might impact immune evasion of PDAC cells — and, in particular, the efficacy of CAR-T cell therapies. Antigen-specific CAR-T cells therapy are approved medicines for several blood cancers^36^; in contrast, treatment of solid tumors, including PDAC, has been met with limited success with cell therapies and there are no approved CAR-T for solid tumors^37^. Although this lack of solid tumor efficacy is multifactorial, PD-L1:PD-1 inhibitory engagement contributes to the limited efficacy of CAR-T therapy^38^ and adding PD-1/PD-L1 inhibitors enhances the efficacy of CAR-T therapy^39^. To test the impact of disruption of PD-L1 regulators on T cell engagement, we isolated primary CD8^+^ and CD4^+^ T cells from three human donors and lentivirally engineered primary human T cells with a CAR that targets mesothelin, a tumor antigen commonly expressed in PDAC^40^ that has been used in multiple clinical trials for PDAC and other solid tumors^41^ (**Fig. 5a**).

**Figure 5.**
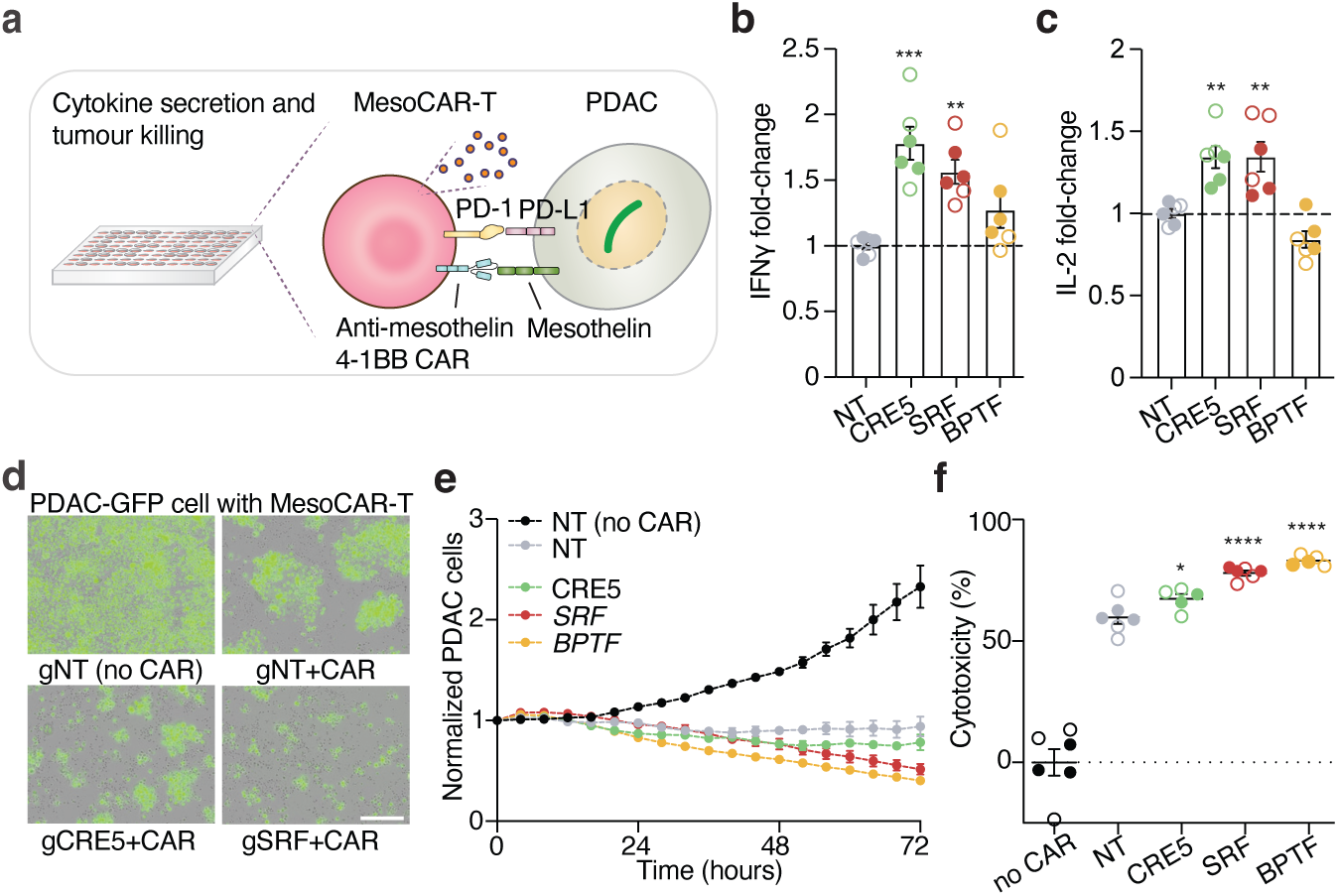
Perturbations on PD-L1 *cis*- and *trans*-regulators improve CAR-T cell response to PDAC. **a,** Anti-mesothelin chimeric antigen receptor T cells (MesoCAR-T) to target pancreatic ductal adenocarcinoma (PDAC) tumors. **b, c,** IFNγ (*b*) and IL-2 (*c)* quantification after 16-hour co-incubation of CD8^+^ MesoCAR-T cells with PDAC BxPC-3 cells transduced with non-targeting (NT), CRE5-targeting, *SRF*-targeting and *BPTF*- targeting gRNAs (*n* = 3 biological replicates per perturbation and donor). Filled and open circles denote two separate human donors. Mann-Whitney *U* test to NT gRNA-transduced cells with Bonferroni-corrected *p*-values : \*\**p* < 0.01, \*\*\**p* < 0.001. **d,** Representative images of BxPC-3-GFP cells transduced with non-targeting (NT), CRE5-targeting or *SRF*-targeting gRNAs co-cultured with human primary CD8^+^ MesoCAR-T cells or untransduced control CD8^+^ T cells. Scale bar, 200 μm. **e,** Counts of BxPC-3-GFP cells transduced with non-targeting (NT), CRE5-targeting, *SRF*-targeting or *BPTF*-targeting gRNAs when co-cultured with human primary CD8^+^ MesoCAR-T cells (*n* = 2-3 biological replicates per donor for 2 donors). Cell counts are normalized to the initial timepoint on a per-well basis and all co-cultures are using a 1:1 effector-to-target cell ratio. **f,** Cytotoxicity of human primary CD8^+^ MesoCAR-T cells when co-cultured with BxPC-3-GFP cells transduced with non-targeting (NT), CRE5-targeting, *SRF*-targeting or *BPTF*-targeting gRNAs after 78 hours at 1:1 effector-to-target cell ratio (*n* = 2-3 biological replicates per donor for 2 donors). Filled and open circles denote two separate human donors. Cytotoxicity is normalized to human primary CD8^+^ cells (no CAR) when co-cultured with BxPC-3-GFP cells transduced with a NT gRNA. Significance was determined using a Mann-Whitney *U* test to NT gRNA-transduced cells with Bonferroni correction: \**p* < 0.05, \*\*\*\**p* < 10^-4^.

To confirm the antigen-specific activity of anti-mesothelin CAR-T (MesoCAR-T), we co-cultured MesoCAR-T cells with two PDAC cell lines (Capan-2 and BxPC-3) that express different levels of mesothelin and a melanoma cell line (A375) lacking mesothelin expression (**Supplementary Fig. 6a**). As expected, MesoCAR-T show antigen-specific tumor killing of Capan-2 and BxPC-3 cells without any activity against the mesothelin-negative A375 cells (**Supplementary Fig. 6b**). To investigate whether the MesoCAR-T model could respond to changes in PD-L1 expression of PDAC cells, we targeted via CRISPR mutagenesis the coding sequence of *PD-L1* and promoter-proximal CRE6 resulted in 86% and 40% loss of PD-L1 expression (**Supplementary Fig. 6c**), respectively, consistent with our prior measurement (**Fig. 1f**). In addition to with the reduced PD-L1 expression, we observed enhanced direct cytotoxicity from MesoCAR-T CD8^+^ cells and increased production of key proinflammatory cytokines IL-2 and IFNγ (**Supplementary Fig. 6d-h**).

Next, we sought to apply this CAR-T model to better understand how the *cis*- and *trans*- regulators identified in the CRISPR screens impact immune evasion of tumor cells. We focused on the SRF-BPTF- CRE5 regulatory module, where we previously identified both *cis*-regulatory elements and TFs that bind at this site. We co-cultured CRE5-, *SRF*- and *BPTF*-disrupted PDAC cells with MesoCAR-T cells from two healthy human donors. Mutagenesis of CRE5 and *SRF* significantly increased IFNγ and IL-2 secretion in MesoCAR-T CD8^+^ cells (**Fig. 5b, c**). Since approved CAR-T therapies in blood cancers use a mixture of CD8^+^ and CD4^+^ T cells, we also tested MesoCAR-T CD4^+^ cells and observed similar increases in IL-2 after targeting CRE5 or *SRF* (**Supplementary Fig. 6i, j**). Most importantly, perturbations of any *cis*- or *trans*- element with the regulatory modulate (CRE5, *SRF* or *BPTF*) led to significant differences in cytoxicity of MesoCAR-T CD8^+^ cells (**Fig. 5d-f**, **Supplementary Fig. 6k-n**). Together, these results demonstrate that genetic disruptions in PD-L1 *cis*- and *trans*-regulators can significantly impact CAR-T cell-mediated immune responses and tumor cell evasion, linking PD-L1 expression level directly to CAR-T cell efficacy in tumor environments.

## The PD-L1 repressor *FOXA3* is recurrently mutated in PDAC tumors and *FOXA3* mutations improve immunotherapy response and survival

We next sought to understand whether our findings regarding *trans*-regulators of PD-L1 expression in human PDAC cells generalize to PDAC tumors from patients. First, we measured if the mutations drive differences in PD-L1 expression, as predicted by the TFome-wide CRISPR screen. To this end, we examined the correlation between PD-L1 *trans-*regulators and *PD-L1* expression in 178 PDAC tumors from The Cancer Genome Atlas. For the activator *trans*-regulators from our CRISPR screen, whose loss resulted in lower PD-L1 expression, we found near-complete concordance. 62 out of the 66 significant TFs in the PD-L1^low^ CRISPR screen, including *STAT1*, *IRF1*, *SRF* and *BPTF*, positively correlate with *PD-L1* expression across the 178 tumors (**Fig. 6a-d**, **Supplementary Table 12**). For repressor *trans*-regulators, we only found a few TFs (*e.g. FOXA3*) where the predicted inverse correlation with *PD-L1* expression was present, which might be explained by the more modest effects overall of repressors (**Supplementary Fig. 7a**).

**Figure 6.**
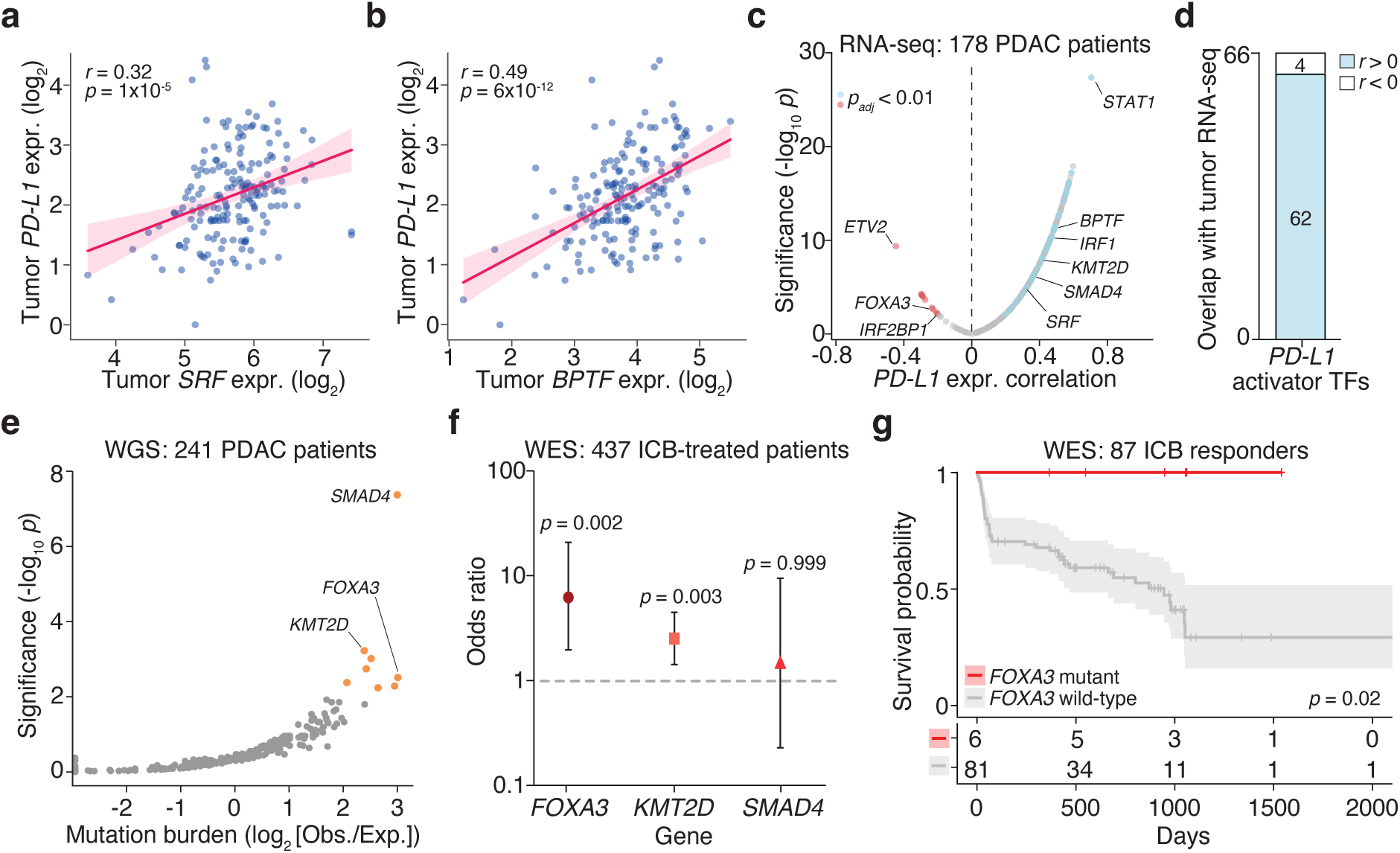
Mutations in PDAC tumors of PD-L1 *trans*-regulators alter *PD-L1* expression and response to immune checkpoint blockade therapies. **a, b,** Expression in primary tumors of *PD-L1* and activator *trans*-regulators *SRF (a)* and *BPTF (b)* (*n* = 178 PDAC patients from The Cancer Genome Atlas^97^). Best fit line with 95% confidence interval. **c,** Pearson correlation between *PD-L1* expression in primary tumors and PD-L1 *trans*-regulators identified in the CRISPR screen (*n* = 178 PDAC patients). TFome-wide screen activator *trans*-regulators are in *blue* and repressor *trans-*regulators are in *red* for significant correlations (*padj* < 0.01). **d**, PD-L1 activator *trans*-regulators identified in the CRISPR screen with concordant or discordant PD-L1 expression *in vivo* (*n* = 178 PDAC patients). **e**, Recurrently mutated PD-L1 *trans*-regulators in 241 PDAC patients from the Pan-Cancer Analysis of Whole Genomes (PCAWG). *Orange* points denote greater than 4-fold enrichment in tumors and FishHook *p* < 0.01. **f,** Response to ICB odds ratio for tumors with high-impact mutations in the indicated gene (two-sided Fisher’s exact test, *n* = 437 patients with exome sequencing and ICB response). Error bars denote the 95% confidence interval. **g,** Overall survival (Kaplan-Meier curve) for patients responding to ICB therapy with (*red*) or without (*gray*) mutations in *FOXA3* (log-rank test, *n* = 87 patients with WES and survival). Shaded areas indicate the 95% confidence interval.

In addition to regulation of *PD-L1*, we sought to understand whether these PD-L1 *trans*-regulators harbor recurrent somatic mutations in PDAC tumors. To this end, we examined 241 PDAC patients with available tumor whole genome sequencing (WGS) from the Pan-Cancer Analysis of Whole Genomes (PCAWG) consortium^42^. For each TF, we modeled the neutral mutational density based on covariates for each gene, including gene size, sequence context, chromatin state, and replication timing^43,44^. Within the PCAWG PDAC cohort, we identified 8 out of 262 PD-L1 *trans*-regulators with significant enrichment of somatic mutations: *SMAD4, KMT2D, ZFP14, EIF5B, FOXA3, GATA6, ZNF296* and *JUNB* (**Fig. 6e**, **Supplementary Table 13**). Among the 8 recurrently mutated genes from the PCAWG WGS cohort, we found 3 *trans-*regulators with concordant changes in *PD-L1* expression (tumor RNA-sequencing): one *PD- L1* repressor (*FOXA3*) and two *PD-L1* activators (*SMAD4, KMT2D*). Taken together, the tumor transcriptomic and WGS datasets suggest that PDAC tumors with mutations in these 3 PD-L1 *trans*- regulators may have altered *PD-L1* expression.

Ultimately, changes in PD-L1 expression are of high clinical import because of the potential to alter response to immune checkpoint blockade (ICB) therapies targeting the PD-1/PD-L1 axis. Tumor expression of PD-L1 correlates with ICB response: PD-L1 expression is a companion diagnostic for PD-1 ICB therapy (c.f. KEYNOTE-001^13,16^) and several recent studies have shown that the effectiveness of ICBs is reduced in the presence of specific mutations^11,28,45^. Therefore, we sought to test whether mutations in these PD-L1 *trans*-regulators might act to reduce the efficacy of immune checkpoint blockade (ICB) therapies.

In order to determine whether recurrent somatic mutations in the three PD-L1 *trans*-regulators may impact ICB response, we assembled 437 patients (from 8 prior studies^11, 45–51^) receiving ICB therapy with tumor whole-exome sequencing (WES) and post-ICB RECIST (Response Evaluation Criteria in Solid Tumors) classifications and/or survival. Using this cohort, we identified mutations in all three *trans*-regulators: *FOXA3* was mutated in 3.4% of patients, *KMT2D* was mutated in 12.2% of patients and *SMAD4* was mutated in 1% of patients. We found that patients with tumor mutations in *FOXA3* and *KMT2D* have higher odds of responding to ICB compared to patients without such mutations (6.2-fold and 2.5-fold, respectively) (**Fig. 6f**, **Supplementary Fig. 7b-d, Supplementary Table 14)**. For *SMAD4*, no significant difference was observed in the response to ICB therapy between patients with and without *SMAD4* mutations (*p* = 0.99). Next, we quantified the association between the mutational status of the *trans*-regulators and overall survival post-ICB treatment. Mutations in *FOXA3* were significantly associated with overall survival in the ICB responders (*n* = 87 ICB responders with WES and survival) (**Fig. 6g**) while *FOXA3* mutations show no effects on survival of ICB non-responders (*n* = 170 ICB non-responders with WES and survival) (**Supplementary Fig. 7e)**, directly linking the *FOXA3*-associated improvements on survival to ICB therapy and its response. Patients with mutations in *KMT2D* had no change in overall survival, despite the difference in ICB response (**Supplementary Fig. 7f)**.

We wondered whether these survival effects were specific to ICB-treated patients or whether mutations in these genes simply modulate survival independent of treatment. We examined survival of patients with or without somatic mutations in *FOXA3*, and *KMT2D* across 10 combined pan-cancer cohorts (*n* = 65,661 patients)^42^. In this much larger cohort, we confirmed that mutations in *FOXA3* did not have any effects on patient survival, suggesting that the impact of *FOXA3* alterations on survival is specific to patients receiving ICB therapy (**Supplementary Fig. 7g**). For *KMT2D*, mutations also had no effect on overall survival (**Supplementary Fig. 7h**). Taken together, these analyses combining PCAWG, TCGA and ICB-treated patient cohorts suggest that TFs that modulate *PD-L1* expression in both human PDAC cells *in vitro* and PDAC patients *in vivo* (such as *FOXA3*) may influence overall survival in ICB-treated patients when mutated.

## Discussion

In this study, we developed paired CRISPR screens to map the *cis*- and *trans*- regulatory landscape of immune checkpoint *PD-L1.* Using these paired CRISPR screens, we identify enhancer and repressor CREs and transcription factors for the expression of *PD-L1* and do so in constitutive and IFNγ-stimulated contexts. Previous studies have probed protein-coding genes that regulate PD-L1 using pooled screens^19,20,52–55^. However, almost 99% of the human genome does not code proteins^9^, and there have been no genetic screens aimed at uncovering noncoding functional elements regulating PD-L1 expression. To decipher the *cis*- regulatory landscape of the noncoding genome, ENCODE and others have profiled DNA accessibility and histone modifications to predict CREs^56^, but, as we have shown, these genetic signatures are not consistently present at CREs. Our saturation mutagenesis CRISPR screen defines CREs at nucleotide resolution and, combined with TFome-wide screen and 3D genome looping, directly identifies the *cis* and *trans* regulatory modules that govern the expression of *PD-L1*.

In most solid tumors, PD-L1 expression is regulated by both IFNγ-dependent and IFNγ-independent pathways^18^. Using H3K27ac-mediated HiChIP, we found that IFNγ reshapes 3D genome conformation globally and in the *PD-L1* TAD. Additionally, the majority of *cis*- and *trans*-regulators identified in our study exhibit context-specific roles that differ between constitutive and IFNγ-stimulated PD-L1 expression. In the constitutive setting, we identified an enhancer regulatory module (CRE-5) bound by transcription factors SRF and BPTF. Perturbations at enhancer-like CRE-5, *SRF* or *BPTF* not only reduce PD-L1 expression but also reduce tumor immune evasion as shown by increased cytotoxicity of MesoCAR-T cells. In addition to their *in vitro* regulation of PD-L1, *SRF* and *BPTF* expression correlates with *PD-L1* expression in PDAC patients *in vivo*. Notably, 62 out of the 66 activator TFs that we identified via perturbations in a PDAC cell model show concordant changes in *PD-L1* expression in patient tumors. We also observed a similar concordance with some negative regulators: The *PD-L1* repressor *FOXA3* identified in our TFome-wide screen is negatively correlated with *PD-L1 in vivo* and harbors recurrent somatic mutations in PDAC patients. Given that PD-L1 expression in of prime clinical importance for PD-1/PD-L1 blockade therapies^16,17^, the positive and negative PD-L1 regulators identified may serve as predictive biomarkers for treatment response: For example, we found that patients harboring mutations in *FOXA3* showed improved checkpoint therapy response and improved overall survival. These predictive biomarkers can used for selecting patients for PD-1/PD-L1 blockade therapies, although larger clinical datasets will be required to validate these findings.

Although ICB has undoubtedly been successful in the clinic^57,58^, recent work suggests that combination strategies can achieve even better outcomes^59,60^. For example, when combined with an inhibitor of FGFR, a well-established cancer driver, the PD-1 inhibitor pembrolizumab led to tumor shrinkage in advanced malignancies^61,62^. Similarly, SHP2 inhibition results in increased PD-L1 and MHC class I expression^63^ and the combination of SHP2 and PD-1^64^/PD-L1^65^ inhibition in mice leads to enhanced immune responses and tumor reduction. An even more germane example is the transcription regulator c-MET, whose inhibition increases PD-L1 expression in multiple solid tumors^66,67^: In phase 3 trials, combination of the PD-1 inhibitor nivolumab and the c-MET inhibitor cabozantinib extended progression-free survival by two-fold in renal cancer, compared to monotherapy^68,69^. In addition to small molecules, combining ICB and CAR-T cells has been shown to enhance the therapeutic activity of CAR-T cells *in vivo* across multiple solid tumors^70,71^ with multiple clinical trials ongoing (NCT04003649, NCT03726515)^72^.

Given the emergence of combination therapies involving the PD-1/PD-L1 axis^39,59^, the novel *cis-* and *trans*- regulators identified here — beyond their roles as predictive biomarkers for ICB response — could also serve as new therapeutic targets in PD-1/PD-L1 combination therapies. More broadly, extensions of this generalizable toolkit can help decipher the regulatory landscape of important genomic loci across diseases and shed light on new therapeutic targets.

## Acknowledgements

We thank the entire Sanjana laboratory for support and advice. We thank the New York University Biology Genomics Core for sequencing resources, E. Mazzoni and O. Qassab for assistance with the CUT&RUN assay, K. Johnson for help with HiChIP analysis, and J. Kang, W. Li and NYU Shanghai for providing laboratory support to X.X. during the COVID-19 pandemic. Z.Z.G. is supported by the National Institutes of Health (NIH) T32 Training Grant (GM136573). N.E.S. is supported by New York University and New York Genome Center funds, NIH/NHGRI (R00HG008171, DP2HG010099, R01HG012790), NIH/NCI (R01CA218668, R01CA279135), NIH/NIAID (R01AI176601), DARPA (D18AP00053), the Melanoma Research Alliance, the Sidney Kimmel Foundation, the MacMillan Center for the Study of the Noncoding Cancer Genome at the New York Genome Center, and the Simons Foundation for Autism Research (Genomics of ASD 896724).

## Author contributions

X.X. and N.E.S. designed the study. X.X. performed the TFome-wide CRISPR screens, functional validations (including arrayed validation and CAR work), AQuA HiChIP experiments and analyzed the data. J.Y.S.K., M.L. and N.B. performed the saturation mutagenesis screen. M.L. designed and cloned the CAR construct. C.M.C. assisted with HiChIP experiments. C.M.C. and Z.Z.G. assisted with HiChIP data analysis. R.E.Y. performed ATAC-seq experiments and assisted with ATAC-seq data analysis. H.W., C.L., X.W. and Z.Z.G. assisted with the TFome-wide screen. X.W. performed CUT&RUN experiments and assisted with CUT&RUN data analysis. Z.Z.G., C.W. and G.G. assisted with PCAWG/TCGA data analysis.

X.X. and N.E.S. wrote the manuscript with input from all authors. N.E.S. supervised the study.

## Additional information

Supplementary Information is available for this paper. Correspondence and requests for materials should be addressed to N.E.S.

## Methods

### Human cell line culture

HEK293FT cells were obtained from ThermoFisher. BxPC-3, A375 and Capan-2 cells were obtained from ATCC. HEK293FT and A375 are cultured in D10: DMEM (Caisson Labs) supplemented with 10% Serum Plus-II (Thermo Fisher). BxPC-3 are cultured in R10: RPMI-1640 (ThermoFisher) supplemented with 10% Serum Plus-II. Capan-2 are cultured in MC10: McCoy’s medium (Caisson Labs) supplemented with 10% Serum Plus-II. Cell lines were regularly tested for mycoplasma and confirmed to be negative using MycoAlert PLUS (Lonza). For IFNγ-stimulated conditions, cells were cultured in medium supplemented with 100 IU/mL human IFNγ (BioLegend) for 48 hours.

### Primary human T cell isolation and cell culture

From the New York Blood Center, we purchased regular buffy coats containing peripheral blood from de-identified healthy donors with informed consent under an IRB-exempt protocol. To isolate the primary human T cells, the peripheral blood mononuclear cells (PBMC) were first isolated from buffy coats using Lymphoprep (Stemcell) gradient centrifugation as previously described^73^. Then, CD8^+^ T cells and CD4^+^ T cells were isolated using the EasySep Human CD8 Positive Selection Kit II (Stemcell) and the EasySep Human CD4^+^ T cell Isolation Kit (Stemcell), respectively. After isolation, both CD8^+^ T cells and CD4^+^ T cells were immediately resuspended in Immunocult-XF T cell Expansion Medium (Stemcell) supplemented with 10 ng/mL recombinant human IL-2 (Stemcell). To activate T cells, we added 25ul Immunocult Human CD3/CD28 T cell Activator (Stemcell) per million T cells per mL. For T cell genome engineering purpose, we transduced primary T cells with lentiviral supernatants 24 h post activation. At 72 h after isolation, engineered T cells were selected with 2 μg/mL puromycin. In the following two weeks, T cells were either split or had the medium (supplemented with 10 ng/mL recombinant human IL-2 and 2 μg/mL puromycin) replaced to maintain a cell density of 1-2 ×10^6^ cells per ml, every 2-3 days. T cells were freshly used for co-culture functional assays between 14 and 21 days after isolation. Alternatively, we cryopreserved engineered T cells between 14 and 21 days after isolation in Bambanker Cell Freezing Medium (Bulldog Bio) and used them for functional assays two days after thawing and recovering them in T cell medium.

### Design and cloning of *cis*- and *trans*-regulatory CRISPR pooled libraries

The *PD-L1* locus saturation mutagenesis gRNA library was designed to tile a 790 kilobase topologically-associated domain (TAD) at the *PD-L1* locus (hg19 chr9: 4,839,758-5,629,082). For these regions, we identified all Cas9- targetable sites containing protospacer adjacent motif (PAM) `NGG` on both strands. The total number of targetable sites identified was 34,204. We eliminated gRNAs with on-target score lower than 0.5^74^, yielding 24,537 gRNAs with a median distance of 12 bp between neighboring gRNAs (**Supplementary Table 3, Supplementary Fig. 1a-c**). In these gRNAs, we have 44 *PD-L1* coding region-targeting gRNAs and 137 *JAK2* coding region-targeting gRNAs as positive controls. For negative controls, we added 1,000 non-targeting gRNAs.

The gRNAs were synthesized as pooled single-stranded oligonucleotides (Twist Biosciences), and then amplified by 10 cycles of PCR using NEBNext High-Fidelity 2X PCR Master Mix (NEB). The resulting amplified DNA were cloned into *Esp3I*-digested (Thermo) lentiCRISPRv2 by Gibson Assembly^75^. The saturation mutagenesis CRISPR library were cloned into the backbone lentiCRISPRv2 (Addgene 52961) containing Cas9, a guide RNA cassette and a puro resistant cassette^75^. Individual gRNAs were cloned into the pCC_01 all-in-one lentiviral vector (Addgene 139086) with an optimized Cas9 scaffold^76^. For the Human TF CRISPR-Cas9 library, we designed gRNAs to target 1,891 known human TFs using the GUIDES web tool (http://guides.sanjanalab.org)^74^ with 10 gRNAs for each TF and added 1,000 non-targeting gRNAs as negative control (**Supplementary Table 7**). The TFome-wide CRISPR library was cloned into the pCC_01 all-in-one lentiviral vector (Addgene 139086) with an optimized Cas9 scaffold^76^.

Cloned libraries were amplified by electroporation into Endura electrocompetent cells (Lucigen). Using dilution plates for colony counting, we verified that all gRNAs were cloned with ≥ 1,000 coverage. For the TFome-wide library, we proportionally (equimolar) spiked in 134 gRNAs, including 124 targeting *PD-L1* CREs identified from the saturation mutagenesis screen, 5 non-targeting gRNAs as negative controls and 5 positive control gRNAs targeting *PD-L1* coding sequences (**Supplementary Fig. 4a, Supplementary Table 7)**.

To verify gRNA representation, we amplified the guide RNA cassette for Illumina sequencing from the pooled library. We performed PCR reactions using 1 ng of the pooled gRNA library plasmids per reaction with Q5 polymerase (NEB). The correct size band from the PCR product was then gel-purified from a 2% E-gel EX (Life Technologies) using the QiaQuick gel extraction kit (QIAGEN). Next, the purified gRNA library was quantified by Tapestation 4200 (Agilent Technologies). Sequencing was performed on the MiSeq instrument using the 150x v3 kit (Illumina).

### Lentiviral production and transduction of pooled libraries

Lentivirus was produced by HEK-293FT cells which were co-transfected with lentiviral packaging plasmid psPAX2^75^ (Addgene 12260), lentiviral enveloping plasmid pMD2.G (Addgene 12259) and gRNA library plasmids at a ratio of 1.0:0.55:0.8, using polyethyleneimine linear MW 25000 (Polysciences). Virus supernatant was harvested 72 h post transfection and filtered through a 0.45 μm Steriflip-HV filter (Millipore). Afterwards, the virus was ultra-centrifuged at 24,000 rpm at 4 C° for 2 h using SW32Ti (Beckman) and the virus pellets were resuspended in RPMI media with 10% FBS to result in a 20-fold concentrated virus and stored at −80 C°.

Before transduction the BxPC-3 cells with concentrated lentivirus in a large-scale for screen, the amount of concentrated lentivirus used for transduction was titrated first to determine the optimum volume of lentivirus delivered into cells to result in 20-30% transduction efficiency. After the lentivirus titration, screen-scale transduction was performed in a format of 12-well cell culture plate, and each well contained 3 x 10^6^ cells with 30 ul lentivirus and 8ug/ml polybrene (Santa Cruz Biotechnology). For the pooled saturation mutagenesis gRNA library, a total of 100 x 10^6^ cells were infected at a multiplicity of infection (MOI) of 0.3. A total of 200 x 10^6^ cells were infected with the TFome-wide gRNA library at a multiplicity of MOI of 0.3 with replicates. We then spun cells at 1,000 G for 2 hours at 30 °C and incubated them overnight at 37 °C. In the following morning, we passaged cells in 225cm^2^ flasks (NEST Scientific). 48 h after transduction, gRNA library transduced cells were selected with 2 ug/mL puromycin, and cells were maintained in media containing 2 ug/mL puromycin. When the puromycin selection completed at day 5 post transduction, we collected ∼25 x 10^6^ cells as an early time point (ETP) control and stored the pellet at −80 °C for gDNA extraction. Throughout the experiment, the representation of each gRNA was monitored (≥ 1000 cells per gRNA).

### PD-L1 pooled CRISPR screens and fluorescence-activated cell sorting

For the pooled saturation mutagenesis screen, we sorted transduced cells into PD-L1^low^ population and PD-L1^high^ population by cell sorting using flow cytometry at day 14. Two days prior to the first sort, 25 x 10^6^ cells were pre-treated with 100 IU/mL IFNγ^19^ and 25 x 10^6^ cells were maintained under constitutive context. On day 14 of the first sorting, cells were collected and counted. An aliquot of 25 x 10^6^ non-treated cells and IFNγ-stimulated cells was frozen down at this step to be used as a presort control. The remaining cells were stained with LIVE/DEAD Violet cell viability dye (ThermoFisher) and anti-PD-L1 antibody (eBioscience). For non-treated cells, PD-L1^low^ cells (corresponding to the bottom 20% of the PD-L1 distribution) and PD-L1^high^ cells (corresponding to the top 20% of the PD-L1 distribution) were collected and cultured one more week for the second round of sorting using the Sony SH800S cell sorter. For IFNγ-stimulated cells, PD-L1^low^ cells and PD-L1^high^ cells were collected and directly frozen down at −80 °C for gDNA extraction. At day 21, non-treated PD-L1^low^ cells and PD-L1^high^ cells were sorted again based on the PD-L1 expression. The sorted cell pellets were stored at −80 °C for gDNA extraction.

For the TFome-wide CRISPR screen, we froze down 20 x 10^6^ cells for each presort control (-IFNγ and +IFNγ) at day 10 post transduction. For IFNγ-stimulated conditions, cells were treated with 100 IU/mL IFNγ for two days. We sorted ∼100 x 10^6^ cells into PD-L1^low^ cells (corresponding to the bottom 10% of the PD-L1 distribution) and PD-L1^high^ cells (corresponding to the top 10% of the PD-L1 distribution) using the Sony SH800S cell sorter for each condition (-IFNγ and +IFNγ).

### Library preparation and sequencing of pooled CRISPR screens

The gDNA was extracted from CRISPR library transduced cells using a modified salting-out precipitation^77^. A two-step PCR protocol was used to amplify the guide RNA cassette for Illumina sequencing from gDNA. For the step1 PCR, we used all gDNA available for each biological sample (10 ug gDNA in each 100 ul reaction using Taq-B polymerase (Enzymatics)). After pooling PCR1 products from multiple reactions, we performed PCR2 reactions (10,000gRNAs per 50ul reaction) using 5 ul of the pooled PCR1 product per reaction with Q5 polymerase (NEB). Each biological sample was amplified with PCR2 primers with unique barcodes (**Supplementary Table 1**). Then we proportionally pooled the PCR2 products of each biological sample and ran the pooled sample through a 2% E-gel EX (Life Technologies), from which we extracted a band of 250-270bp and purified it with the QiaQuick gel extraction kit (QIAGEN). Next, the purified sample was quantified by Tapestation 4200 (Agilent Technologies). Sequencing was performed on the NextSeq 500-II machine using the MidOutput mode with the 150x v2.5 kit (Illumina).

To analyze the sequencing data, we first demultiplexed the samples using bcl2fastq (Illumina), trimmed off the 5’ adaptor for each biological sample with Cutadapt v.1.13 (`-m 24 -e 0.1 --discard-untrimmed`)^78^ and aligned the gRNA sequences to the reference CRISPR library using Bowtie v.1.1.2 (`-v 1 -m 1 --best -- strata`)^79^. The gRNA counts in each biological sample were normalized to the total number of reads and log2- transformed. To calculate gRNA enrichment score, we measure the fold-change of each gRNA between the sorted sample (e.g. PD-L1^low^ -IFNγ, PD-L1^high^ -IFNγ, PD-L1^low^ +IFNγ and PD-L1^high^ +IFNγ) and the corresponding inputs (e.g. presort -IFNγ and presort +IFNγ).

### *Cis*- and *trans*-regulatory CRISPR screen data analysis

In the saturation mutagenesis screen, we have 24,537 targeting gRNAs with a median distance of 12 bp between neighboring gRNAs. To identify the candidate *cis*-regulatory elements for PD-L1 expression, we first developed a rolling window-based analysis where each window consists of five consecutive gRNAs with a medium size of 100bp (**Supplementary Fig. 2b**). After ranking the enrichment of 5 gRNAs in each window, we select the second highest value to represent the enrichment score of the window. To identify the significant enriched window, we randomly selected 5 gRNAs from the gRNA library (∼25,537) and used the second highest enrichment score to represent the simulated window performance using the permutation test (*n* = 250,000)^22^. Compared to the simulated windows, we identified the candidate CREs from the 24,533 5-gRNA windows with *p* <10^-3^.

For the TFome-wide CRISPR screen, we performed the enrichment analyses at both the gRNA and TF level using FDR-adjusted robust ranking algorithm (RRA) *p*-values. We use 20 gRNAs targeting the same TF in two replicates for the RRA. To define candidate *trans*-regulators for PD-L1 expression, we selected enriched TFs with FDR-adjusted robust ranking algorithm (RRA) *p*-value <0.01.

### Array validation of candidate *cis*-regulators and *trans*-regulators

To validate PD-L1 cCREs, we first selected 43 CREs distributed across four sorted cell populations (include two CREs-CRE6 and CRE7-located at *PD-L1* promoter) and picked two gRNAs with the highest and the second highest enrichment by the normalized log2(sorted/presort) count ratio of each 5-gRNA targeting CRE. For PD-L1 *trans*- regulators, we selected 3 activator TFs and 2 repressor TFs across four sorted cell populations and used 4 gRNAs (two from the pooled TFome-wide screen and two newly designed using the GUIDES web tool (http://guides.sanjanalab.org)^74^).We also included two *PD-L1* coding sequence-targeting gRNAs and 3 non-targeting sgRNAs to serve as positive and negative controls, respectively (**Supplementary Table 1**). For individual gRNA, standard desalted short oligonucleotides (IDT) were annealed and cloned into the pCC_01 (Addgene 139086)^76^. After producing lentivirus containing the individual gRNA, BxPC-3 cells were transduced with lentiviral supernatants. After puromycin selection and expansion for 7 days, cells were treated with or without 100 IU/mL IFNγ^19^ (BioLegend) for 2 more days.

At day 9, cells were processed to measure PD-L1 expression via flow cytometry. First, the cells were collected, washed once with D-PBS and stained with LIVE/ DEAD Violet cell viability dye (ThermoFisher) for 5 minutes at room temperature in the dark, followed by additional staining with APC-conjugated anti-PD-L1 antibody for 20 minutes on ice. After PD-L1 antibody staining, the cells were washed with PBS three times and acquired on a BD-Aria II machine for the saturation mutagenesis screen and a Sony SH800S cell sorter for the gRNA array validation and the TFome- wide screen. Flow cytometry data was analyzed using FlowJo (Treestar).

### Anti-mesothelin CAR-T cell engineering

We used the lentiviral backbone from lentiCRISPRv2 (Addgene 52961)^75^. Anti-mesothelin (SS1) and 4-1BB (BBz) CAR^80,81^ were synthesized as gBlocks (IDT) (**Supplementary Table 1**). After adding Gibson overhangs by PCR (**Supplementary Table 1**), anti-Mesothelin (SS1) and 4-1BB (BBz) CAR were inserted into *Xba*I- and *BamHI*-cut lentiCRISPRv2 to generate pOT_7 (lenti-EFS-SS1-BBz-2A-puro, Addgene 217990). After producing lentivirus expressing lentiCAR plasmid, we concentrated lentiviral supernatants using Lentivirus Precipitation Solution (Alstem) and transduced primary T cells at 24 hours post isolation. The engineered CAR-T cells were used for functional assays after 14 days.

### CAR-T cell cytokine secretion assays

To determine the effects of CRE knockout on CAR-T associated tumor immune evasion, we co-cultured CRE- disrupted PDAC cells (BxPC-3 and Capan-2) and CAR-T cells at 1:2 effector:target (E:T) ratios to detect the IL-2 or IFNγ secretion . First, primary human T cells (both CD8^+^ and CD4^+^) were isolated from human PBMC and transduced with lentiviral vector encoding anti-mesothelin CAR. Untransduced primary CD8^+^ and CD4^+^ T cells were maintained in parallel with CAR-T cells as negative controls.

For PDAC cells, we mutagenized CREs using CRISPR CRE-targeting gRNAs. PDAC cells transduced with non-targeting gRNAs were included as negative controls. When both CAR-T cells and PDAC cells were lentivirally transduced and ready to be used, T cells were rested for 24 hours in Immunocult-XF T cell Expansion Medium (Stemcell) without IL-2. After 24 hours, both T cells and PDAC cells were counted, plated at 1:1, 1:2, 1:4 and 1:8 effector:target (E:T) ratios (5 × 10^4^ PDAC cells and 2.5 × 10^4^ T cells at 1:1 E:T ratio) per well in triplicate in a round bottom 96-well cell culture plate and cultured in Immunocult-XF medium without IL-2 for 16 hours. After centrifuging the culture plate at 700g for 5 minutes, cell supernatant in each well were collected, diluted 3 times with Immunocult-XF medium without IL-2 and used for cytokine quantification with Human IL-2 or IFNγ DuoSet (R&D Systems) enzyme-linked immunosorbent assay and read out using an Infinite F200 Pro (Tecan) plate reader.

### CAR-T cell cytotoxicity assay

PDAC cells (BxPC-3 and Capan-2) cells were first transduced with a lentiviral vector encoding EGFPd2PEST-NLS and a hygromycin resistance gene. The PDAC GFP cells were kept in hygromycin selection throughout the culture, to maintain stable EGFP expression, and hygromycin was only removed from the medium before the co-culture assay. T cells and PDAC cells were engineered using lentivirus to introduce anti-mesothelin CAR and knockout PD-L1 CREs, respectively. Next, T cells were rested for 24 hours in medium without IL-2. We co-cultured PDAC cells and CAR-T cells at 1:1, 1:2, 1:4 and 1:8 E:T ratios in triplicate in a flat 96-well plate. The cells were incubated for 2h at 37 C° to allow the cells to fully settle down after plating them. The plates were then imaged using an Incucyte SX1, using 20× magnification and acquiring 4 images per well every 2 hours. For each well, the integrated GFP intensity was normalized to the initial time point (0 hours / initial time point).

### H3K27ac AQuA-HiChIP sample preparation and analysis

We prepared HiChIP cell libraries for H3K27ac AQuA-HiChIP^26^. We cultured BxPC-3 cells in R10 only or in R10 supplemented with 100 IU/mL IFNγ (BioLegend) for 2 days (*n* = 6 biological replicates per condition). For absolute quantification (AQuA-HiChIP), we grew NIH3T3 mouse fibroblasts in DMEM media (supplemented with 10% Serum Plus-II (Thermo Fisher Scientific). Then, we fixed cells in 1% formaldehyde (Sigma-Aldrich) for 10 mins and quenched to a final concentration of 125nM glycine (Sigma-Aldrich). We spiked in 2 million fixed mouse cells with 10 million of fixed treated-BxPC-3 cells or non-treated cells. The cells were lysed in 0.5% SDS (Sigma-Aldrich), quenched with 10% Triton X-100 (Sigma-Aldrich), and digested with *MboI* (NEB R0147M). Next, the DNA overhangs were blunted, biotinylated (ThermoFisher 19524016), and ligated. Nuclei were spun down, resuspended in nuclear lysis buffer and sonicated using a Covaris LE220 program: Fill level 10, PIP 450, Duty factor 30, CPB 200. We incubated the sheared DNA with Dynabeads Protein A (ThermoFisher 10001D) for 2 hours at 4 °C to preclear the samples. We then placed the tubes on a magnet and the supernatant was kept. We performed immunoprecipitation with a cross-species reactive H3K27ac antibody (Active Motif 39133). The samples were incubated with the antibody overnight at 4 °C. Next, we added Dynabeads Protein A into tubes post overnight incubation for 2 hour-incubation at 4 °C. We then washed, eluted and treated the samples with Proteinase K (New England Biolabs P8107S). We purified the samples using Zymo ChIP DNA Clean & Concentrator. Biotin capture was performed with Dynabeads M-280 Streptavidin (ThermoFisher 11205D), followed by library preparation^26^. We purified the amplified libraries with Illumina Sample Purification Beads. We sequenced the libraries using paired-end reads with NovaSeq 6000 S4 2x150 kit to generate >200 million read pairs per sample.

We mapped HiChIP paired-end reads to hg38 (human) and mm10 (mouse) using HiC-Pro^82^. Default settings were used to remove duplicate reads and identify unique valid contacts. After measuring the AQuA factor^26^, the valid contacts of 6 replicates across conditions are merged to map the contacts and call HiChIP loops. To call HiChIP loops, unique valid contacts were binned at 5 kb or 2kb resolution and filtered for loops that had 3 or more valid contacts in one bin and were with loop length equal or larger than the bin size. The HiChIP loop strength (count of valid contacts between bins) is normalized by AQuA factor in the HiChIP samples (**Supplementary Table 6**). When quantifying the promoter-anchored or enhancer-anchored loops, the 5kb-anchors of HiChIP loops were intersected with the promoter of genes (TSS±1kb) or PD-L1 CREs by bedtools v2.25.0. Given the promoter of *PLGRKT* and *PD-L1* reside in two neighboring bins, these loops are shown as *PD-L1* contacts.

### ATAC-seq sample preparation and analysis

ATAC-seq was performed^83^ using homemade transposomes^84^ with minor modifications that are detailed below. For each replicate, we trypsinized, washed, and resuspended 250,000 cells in lysis buffer (10 mM Tris-HCl, pH 7.4, 10 mM NaCl, 3 mM MgCl2 and 0.1% NP-40, 0.1% tween-20, 0.01% digitonin)^85^ and lysed for 10 minutes on ice. After lysis, 50,000 cells were resuspended in transposase reaction mix (1ul Tn5 (0.5 µM), 4ul 5X TD-TAPS (50mM TAPS-NaOH, 25mM MgCl2, 50% DMF), 5ul dilution buffer (10mM Tris-HCl, pH 7.4, 100mM NaCl, 1mM DTT, 50% glycerol), 2ul 10% Tween, and 8ul PBS) and tagmentation was performed at 37°C for 30 minutes. Next, samples were purified using the Zymo Clean and Concentrate kit and PCR amplified with NEBNext High-Fidelity 2× PCR Master Mix. The PCR program was as followed: 72 °C for 5 minutes, 98 °C for 30 seconds, (98 °C for 10 seconds, 63 °C for 30 seconds, 72 °C for 60 seconds) x 12 cycles. Samples were sequenced on both Illumina NextSeq 500 and Ultima Genomics platforms.

For Illumina data, adaptors were trimmed with trimmomatic (version 0.39)^86^ with the options `ILLUMINACLIP:Trimmomatic-0.39/adapters/NexteraPE-PE.fa:2:30:10 LEADING:3 TRAILING:3 SLIDINGWINDOW:4:25 MINLEN:24`. For Ultima data, cram files were converted to fastq using samtools (version 1.9)^87^, and trimmed using cutadapt (version 4.0)^78^. Reads from both sequencing platforms were aligned to GRCh38 with bowtie2 (version 2.4.4)^79^, deduplicated with picard through gatk MarkDuplicates (version 4.2.5.0), and peaks were called with MACS2 callpeak (version 2.2.7.1)^88^ with default parameters.

### Integration of *cis*-regulators and *trans*-regulators

To pair PD-L1 *trans*-regulators to specific PD-L1 CREs, we first identified the significantly enriched TF motif(s) at each CRE. TF motif prediction was carried out by extending 100 bp of genomic sequence centered on each end of the PD-L1 CRE. The sequences were entered into the two TF motif profile databases - JASPAR with motifs of 779 TFs^33^ and HOCOMOCO with motifs of 678 TFs^34^, respectively. FIMO tool was then used to identify high-confidence enriched motif(s) (*p* ≤ 10^-4^) by position-weight matrix matching at each CRE^89^. Integrating the significance of PD-L1 *trans*-regulators in our TFome-wide CRISPR screen with the enrichment of TF motifs in our *cis*-regulators, we identified the candidate CRE-TF interaction modules for PD-L1 regulation with CRISPR TF (RRA) *p* ≤ 10^-2^ and the enriched motif significance *p* ≤ 10^-4^.

### CUT&RUN sample preparation and analysis

CUT&RUN was performed in triplicate for each target protein using BxPC-3 cells with CUTANA ChIC/CUT&RUN Kit (EpiCypher)^90^. For each replicate, 500,000 cells were trypsinized, washed, and bound to activated Concanavalin A-conjugated paramagnetic beads (EpiCypher). The cell-bead conjugates were then resuspended in recommended buffer with 0.5 μg antibody targeting the protein of interest: Anti-BPTF (ThermoFisher Scientific A300-973A), Anti-SRF (Cell Signaling Technology 5147S), Anti-H3K4me3 (EpiCypher 13-0041k) as positive control, and Anti-IgG (EpiCypher 13-0042k) as negative control. The suspensions were incubated overnight at 4 °C with gentle shaking. Then, the cell-bead conjugates were washed twice with the kit’s permeabilization buffer and incubated with pAG- MNase (Epicypher 15-1016) for 10 min at room temperature. Subsequently, the unbound pAG-MNase was washed away, and the MNase was activated by adding CaCl2 to a final concentration of 2 mM. The reaction was stopped after the pAG-MNase was allowed to digest and release the antibody-bound chromatin for 2 hours at 4 °C. Finally, the released DNA was purified with the spin columns provided by the kit (EpiCypher). Library preparation was performed using NEBNext Ultra II DNA Library Prep Kit (New England Biolabs). The libraries were sequenced on a Illumina NextSeq 500 using 2x75 bp reads.

The fastq files were trimmed and quality-controlled with Trim Galore (version 0.6.10 with the default options in a pair-end mode. The resulting reads were aligned to hg38 using bowtie2 (version 2.4.4)^79^ with options `--local -- very-sensitive-local --no-unal --no-mixed --no-discordant --dovetail -I 10 -X 700`. The alignment files were sorted by coordinate, then unpaired reads were removed, and indexed with samtools (version 1.14)^87^. For reads visualization on genome tracks, the processed bam files were merged with samtools and visualized via the Integrative Genomics Viewer (version 2.16.1)^91^. For peak identification, peaks and summits were called with MACS2 (version 2.2.7.1)^88^ using all three individual bam files for all replicates targeting the protein of interest (`-t`) and all three targeting IgG as controls (`-c`) with options `--format BAMPE --gsize 2.7e9 --keep-dup all`. The ENCODE blacklist v2 for hg38^92^ was included in computeMatrix as `--blackListFileName` input to filter out reads aligning to problematic genome regions. For making the heatmap of reads on peak regions, deepTools (version 3.5.0)^93^ was used to create the read score matrices with the function `computeMatrix reference-point --referencePoint center - -upstream 2000 --downstream 2000 --sortRegions descend`.

### HiC-seq, RNA-seq and ChIP-seq analyses

HiC data of BxPC-3 was downloaded from NCBI Gene Expression Omnibus (GEO) (accession GSM5605646^94^) and visualized in The 3D Genome Browser^95^. H3K27ac ChIP-seq data of BxPC-3 was downloaded from NCBI GEO as processed bigWigs (accession GSM3178671^96^). These data were visualized and integrated in IGV. The RNA-seq data of BxPC-3, Capan-2 and A375 was from the CCLE in DepMap portal as normalized log2(TPM+1) count.

### *In vivo* expression of PD-L1 and its *trans*-regulators

Gene expression quantification data (derived from bulk RNA-Seq) was downloaded for 178 PDAC samples (Project ID: TCGA-PAAD, https://portal.gdc.cancer.gov/projects/TCGA-PAAD) from the National Cancer Institute’s Genomic Data Commons^97^. For all samples, TPM (transcripts per million) values were extracted for PD-L1 and for each of the significant PD-L1 *trans*-regulators from the TFome-wide CRIPSR screens. Pearson correlation coefficients were calculated between the log2($+1)-transformed TPM of *PD-L1* and each of PD-L1 *trans*-regulators. We corrected *p-*values using the Bonferroni method, determining significant correlations as those with *padj* < 0.05.

### Identification of recurrently mutated PD-L1 *trans*-regulators under positive selection

We identified genes under positive selection using the fishHook statistical framework (https://github.com/mskilab/fishHook)^43^, which has also been used in prior studies^44^. Briefly, we used fishHook to fit a gamma-Poisson model of neutral mutational density across the genome with replication timing, chromatin state, and sequence context as covariates to correct for their effects on mutational density. We then compared the observed mutational rates in the context of the given covariates in the PD-L1 *trans*-regulators to the neutral model to determine significance.

Specifically, we fit the model using whole genome sequencing mutations from 241 PDAC patients across 19,688 GENCODE genes^98^ that had metadata on GeneCards^99^, excluding any mutations that did not fall within exons. For covariates we used a pan-cancer covariate set (“covariome”) from our previous work^44^ that included principal components of replication timing across 96 cell lines, ChromHMM states across 127 cell lines, and mono-, di-, and tri-nucleotide context across the genome. We then compared the mutational rates in the PD-L1 *trans*-regulators in the context of the covariates to assess for positive selection and nominated genes using a threshold of fishhook^100^ *p* < 0.01 and log2(enrichment of mutation count) > 2.

### Overall survival of immune checkpoint blockade-treated and pan-cancer patients

We utilized a cohort of cancer patients treated with ICB therapy who had matched WES derived from 8 studies (*n* = 437 patients)^11,45–51^ that were previously harmonized^44^. Genes were classified as mutated in a patient if they had one or more high or moderate impact variants (i.e. non-synonymous variants) within a given gene. For patients with overall survival data (*n* = 216 patients), the difference in overall survival between the altered and non-altered groups was calculated using a two-sided log-rank test. For our pan-cancer analysis, we downloaded the overall survival data of patients with or without somatic mutation in *SMAD4*, *FOXA3*, and *KMT2D* across 10 combined pan-cancer cohorts from the cBioPortal (*n* = 35,403 patients, *n* = 531 patients, and *n* = 35,390 patients, respectively)^42,45,101–108^. The difference in overall survival between the altered and non-altered groups was calculated using a two-sided log-rank test.

## Data availability

H3K27ac AQuA-HiChIP, ATAC-seq and CUT&RUN datasets have been deposited in NCBI’s Gene Expression Omnibus repository^109,110^ under GEO Series accession number GSE255701.

