## Supplementary Figures for "Paired CRISPR screens to map gene regulation in *cis* and *trans*"


**Supplementary Fig. 1.** Design and analysis of CRISPR-Cas9 saturation mutagenesis screen for PD-L1 *cis*-regulatory elements.

**Supplementary Fig. 2.** Identification and validation of candidate *cis*-regulatory elements (cCREs) for PD-L1 expression.

**Supplementary Fig. 3.** Quantification of H3K27ac HiChIP contacts in untreated and IFNγ-treated BxPC-3 cells.

**Supplementary Fig. 4.** TFome-wide CRISPR screen for PD-L1 *trans*-regulators.

**Supplementary Fig. 5.** Integration of PD-L1 *cis*-regulatory elements and *trans*-regulators.

**Supplementary Fig. 6.** PD-L1 *cis*- and *trans*-regulators impact CAR-T mediated immune evasion.

**Supplementary Fig. 7.** Characterization of PD-L1 *trans*-regulators in PDAC patients and immune checkpoint blockade (ICB)-treated patients.


**Supplementary Tables**


**Supplementary Table 1.** Oligonucleotide sequences for Illumina sequencing primers, CAR-T cloning primers and CRISPR gRNAs in arrayed TF validation.

**Supplementary Table 2.** Antibodies for flow cytometry and CUT&RUN.

**Supplementary Table 3.** *Cis*-regulatory element CRISPR mutagenesis guide RNA library.

**Supplementary Table 4.** Rolling window analysis of candidate *cis*-regulatory elements.

**Supplementary Table 5.** Candidate *cis*-regulatory element PD-L1 protein expression.

**Supplementary Table 6.** H3K27ac AQuA-HiChIP quantitation.

**Supplementary Table 7.** TFome-wide library targeting ~2,000 human TFs.

**Supplementary Table 8.** Fold enrichment and validation of TFs in the *trans*-regulator screens and RNA expression of TFs from CCLE.

**Supplementary Table 9.** TF motif enrichment in the candidate *cis*-regulatory elements.

**Supplementary Table 10.** TF motifs and CRE-targeting gRNAs for the 25 predicted TF-CRE regulatory modules.

**Supplementary Table 11.** Genome-wide binding sites of SRF and BPTF from the CUT&RUN assay.

**Supplementary Table 12.** Correlation between *trans*-regulators and *PD-L1* of RNA expression in PDAC tumors.

**Supplementary Table 13.** Somatic mutations of *trans*-regulators in PDAC patients.

**Supplementary Table 14.** Somatic mutation, ICB-response and survival of ICB-treated patients.

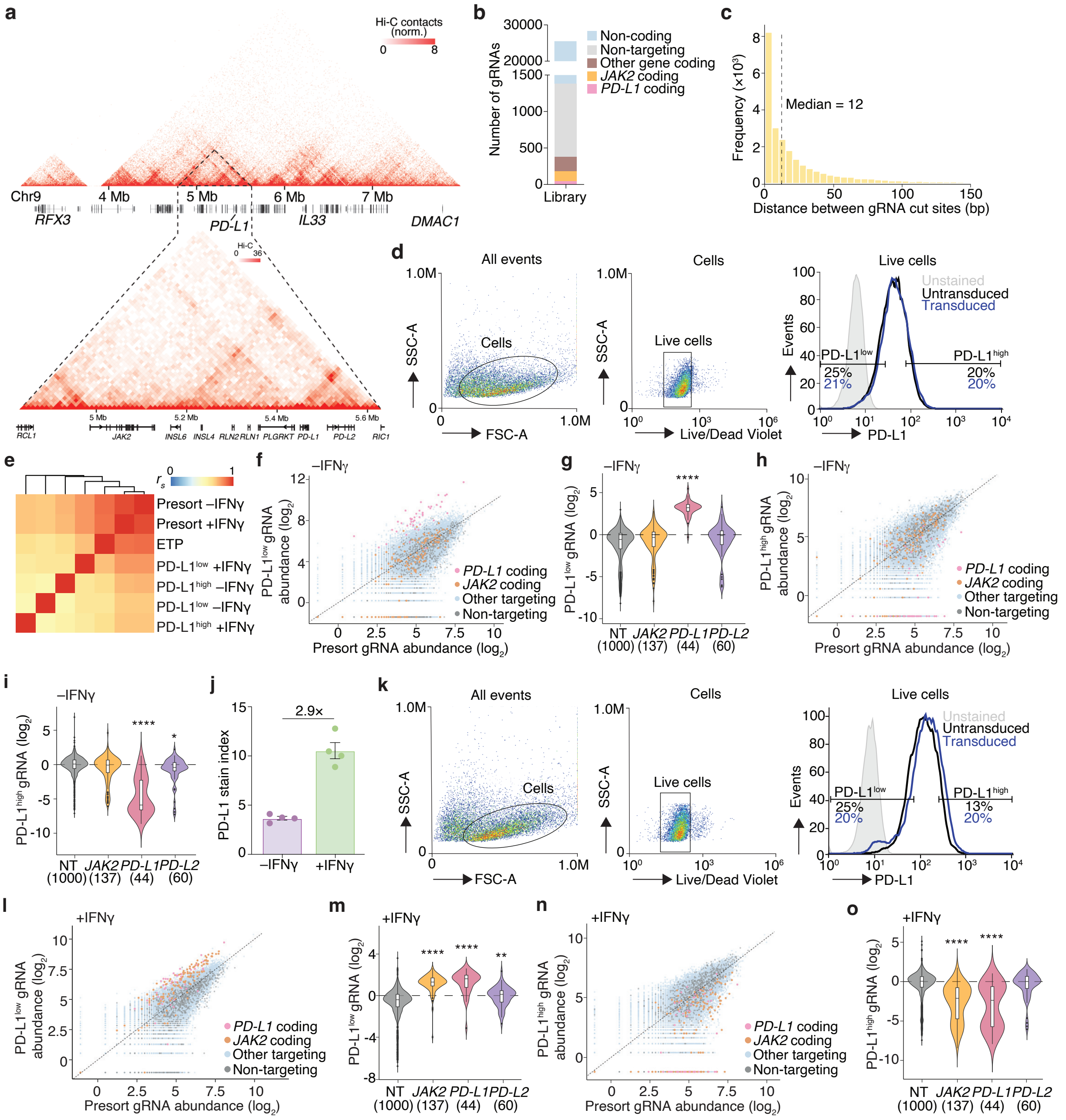

**Supplementary Fig. 1 | Design and analysis of CRISPR-Cas9 saturation mutagenesis screen for PD-L1 *cis*-regulatory elements.**

**a,** HiC contact mapping around *PD-L1*. The *PD-L1* topologically associating domain (TAD) in BxPC-3 cells is shown (enlarged region).

**b,** Target sites of 25,537 gRNAs in the CRISPR saturation mutagenesis library tiling across *PD-L1* locus.

**c,** Distances between the cut sites of neighboring gRNAs in the saturation mutagenesis screen library.

**d,** Sequential gating strategy for PD-L1 expression in untreated cells.

**e,** Spearman correlations between normalized gRNA counts of the saturation mutagenesis CRISPR library-transduced BxPC-3 cells.

**f,** Normalized read counts for each CRISPR gRNA in cell populations before sorting and after PD-L1$^{low}$ sorting (without IFNγ stimulation).
**g,** Fold-change of gRNAs targeting the coding sequences (CDS) of *PD-L1, JAK2* and *PD-L2* in PD-L1$^{low}$ cell populations compared with presort cell populations (without IFNγ stimulation).
**h,** Normalized read counts for each CRISPR gRNA in cell populations before sorting and after PD-L1$^{high}$ sorting (without IFNγ stimulation).
**i,** Fold-change of gRNAs targeting the CDS of *PD-L1, JAK2* and *PD-L2* in PD-L1$^{high}$ cell populations compared with presort cell populations (without IFNγ stimulation).
**j,** Stain index of constitutive and IFNγ-stimulated expression of PD-L1 ($n = 4$ biological replicates).
**k,** Sequential gating strategy for PD-L1 expression in IFNγ-treated cells.
**l,** Normalized read counts for each CRISPR gRNA in cell populations before sorting and after PD-L1$^{low}$ sorting (with IFNγ stimulation).
**m,** Fold-change of gRNAs targeting the CDS of *PD-L1, JAK2* and *PD-L2* in PD-L1$^{low}$ cell populations compared with presort cell populations (with IFNγ stimulation).
**n,** Normalized read counts for each CRISPR gRNA in cell populations before sorting and after PD-L1$^{high}$ sorting (with IFNγ stimulation).
**o,** Fold-change of gRNAs targeting the coding domain sequences (CDS) of *PD-L1, JAK2* and *PD-L2* in PD-L1$^{high}$ cell populations compared with presort cell populations (with IFNγ stimulation).
For all panels, unless otherwise specified, significance was determined using a Mann-Whitney *U* test to the non-targeting (NT) group with Benjamini-Hochberg correction: $*p < 0.05$, $**p < 0.01$, $***p < 0.001$ and $****p < 0.001$.

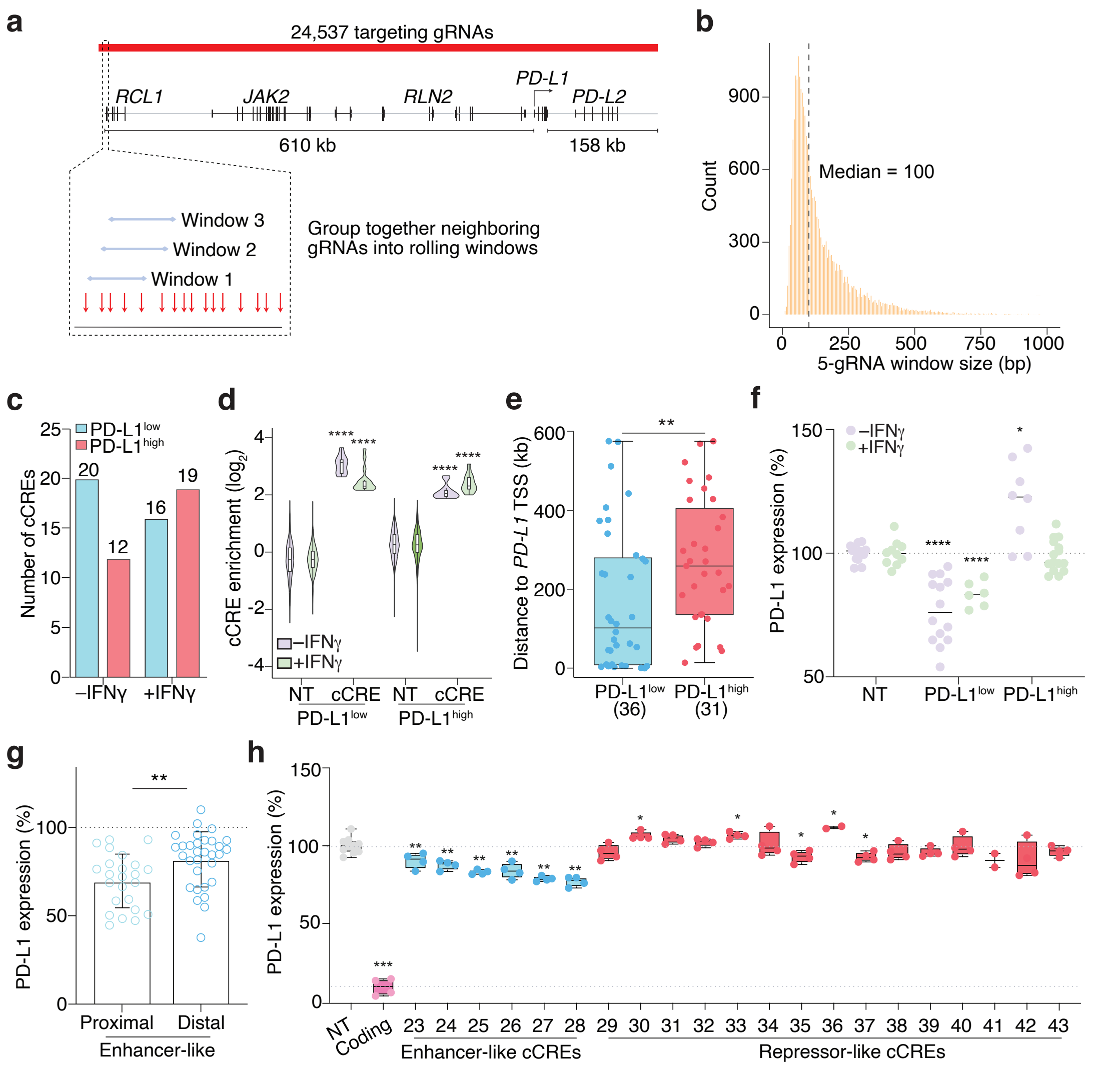


**Supplementary Fig. 2 | Identification and validation of candidate *cis*-regulatory elements (cCREs) for PD-L1 expression.**

**a,** Rolling window analysis for the identification of PD-L1 cCREs.

**b,** Lengths of rolling windows analyzed (five gRNA span) from the saturation mutagenesis screen.

**c,** Enriched cCREs in the four sorted populations.

**d,** Enrichment (fold-change normalized to non-targeting [NT] gRNAs) of cCRE-targeting gRNAs in untreated cells and IFNγ-treated cells. There are 1000 NT gRNAs and 5 gRNAs for each cCREs in the four sorted populations.

**e,** Distance between enhancer-like cCREs (enriched in PD-L1$^{low}$ cell populations) and repressor-like cCREs (enriched in PD-L1$^{high}$ cell populations) to the transcription start site (TSS) of *PD-L1* in untreated cells.

**f,** PD-L1 expression after individual Cas9 targeting (mutagenesis) of enhancer-like (enriched in PD-L1$^{low}$) and repressor-like (enriched in PD-L1$^{high}$) cCREs in untreated and IFNγ-treated cells. Dots in PD-L1$^{low}$ and PD-L1$^{high}$ groups denote the mean change of PD-L1 expression ($n$ = 2 gRNAs with 2 biological replicates per cCRE).

**g,** PD-L1 expression after individual Cas9 targeting (mutagenesis) of proximal enhancer-like cCREs (cCREs 5 – 10) and distal enhancer-like cCREs in untreated cells (*n* = 2 gRNAs with 2 biological replicates per cCRE).

**h,** IFNγ-stimulated PD-L1 expression after transduction with NT, *PD-L1* coding exon-targeting (Coding), enhancer-like and repressor-like cCRE-targeting gRNAs (*n* = 1 - 3 gRNAs with 2 - 4 biological replicates). Significance was determined by Bonferroni-corrected Mann-Whitney *U* test.

For all panels, significance was determined by the Mann-Whitney *U* test unless otherwise specified and $*p < 0.05$, $**p < 0.01$, $***p < 0.001$, $****p < 0.001$.

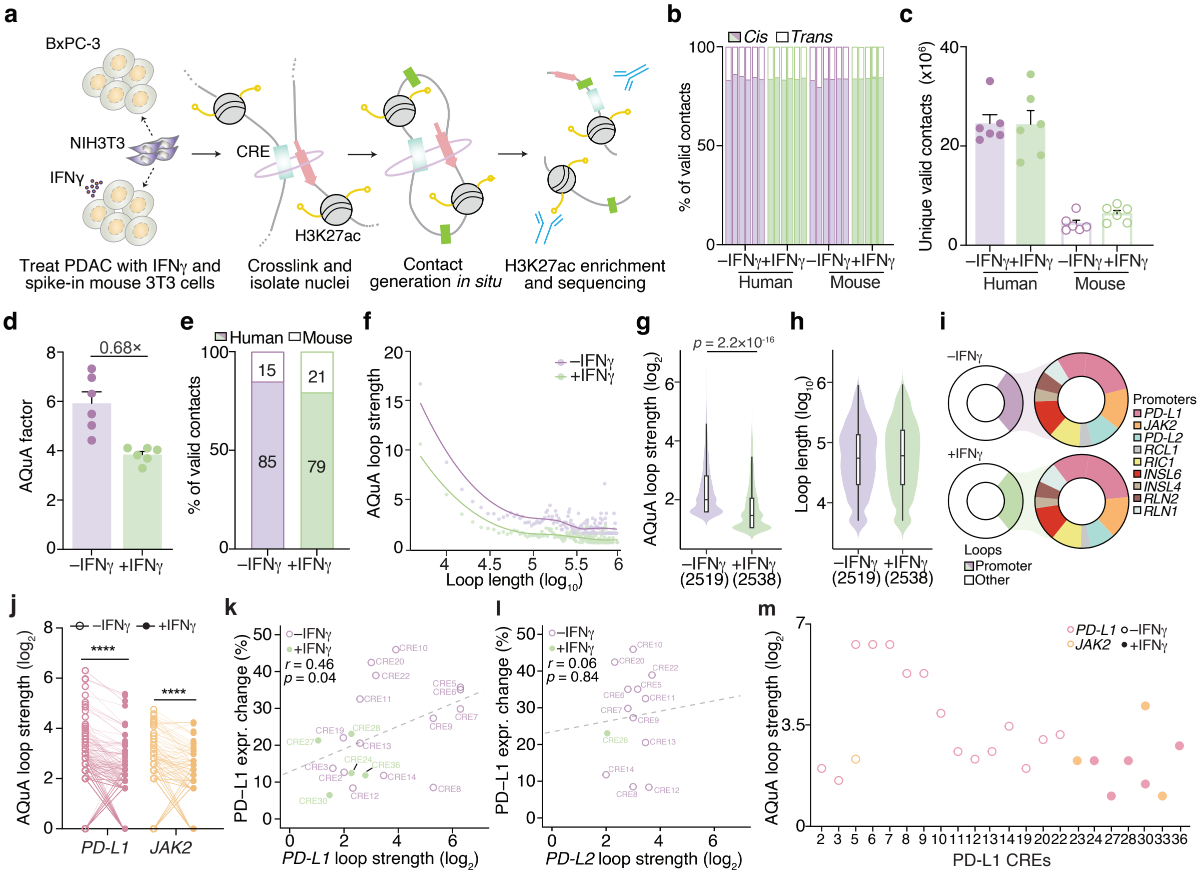


**Supplementary Fig. 3 | Quantification of H3K27ac HiChIP contacts in untreated and IFNγ-treated BxPC-3 cells.**

**a,** Absolute quantification of chromatin architecture (AQuA) H3K27ac HiChIP in untreated and IFNγ-treated BxPC-3 cells. For absolute quantification, each condition received the same mouse spike-in (NIH3T3 cells).

**b**, *Trans* (inter-chromosome) and *cis* (intra-chromosome) contact frequencies for AQuA H3K27ac HiChIP among samples experiments ($n = 6$ biological replicates).

**c,** Valid unique H3K27ac HiChIP contacts across species in untreated and IFNγ-treated BxPC-3 cells.

**d,** AQuA scaling factor for AQuA-HiChIP in untreated and IFNγ-treated BxPC-3 cells.

**e,** The comparison of overall mouse to human H3K27ac HiChIP contacts in untreated and IFNγ-treated BxPC-3 cells.

**f,** AQuA-normalized H3K27ac HiChIP loop strength (AQuA-normalized count of valid contacts at 5 kb resolution) as a function of loop lengths (distance between anchors) within the *PD-L1* TAD (hg38, chr9: 4,775,560 – 5,780,418) in untreated and IFNγ-treated BxPC-3 cells. The average strength of loops with the same length is plotted.

**g**, AQuA-normalized H3K27ac HiChIP loop strength in untreated and IFNγ-treated BxPC-3 cells. Numbers in parentheses indicate the number of loops in the *PD-L1* TAD for each condition. Significance was determined by the Mann-Whitney *U* test.

**h**, H3K27ac HiChIP loop length (bp) in untreated and IFNγ-treated BxPC-3 cells. Numbers in parentheses indicate the number of loops in the *PD-L1* TAD for each condition.

**i**, Promoter-anchored H3K27ac HiChIP loops in the *PD-L1* TAD.

**j,** Change (between constitutive and IFNγ-stimulated contexts) in AQuA-normalized H3K27ac HiChIP loop strength for *PD-L1* or *JAK2* promoter-anchored loops in the *PD-L1* TAD. Significance was determined by the Wilcoxon matched-pairs signed rank test.
**k, l**, Percent change in PD-L1 protein expression after CRISPR perturbation for the indicated CREs and the AQuA-normalized H3K27ac HiChIP loop strength for loops between the indicated PD-L1 screen CRE and the *PD-L1* (*k*) or *PD-L2* (*l*) promoter ($n$ = 2 biological replicates of 2 gRNAs per CRE).
**m,** AQuA-normalized H3K27ac HiChIP loop strength for loops between the indicated PD-L1 screen CRE and the *PD-L1* (*pink*) or *JAK2* (*orange*) promoter.

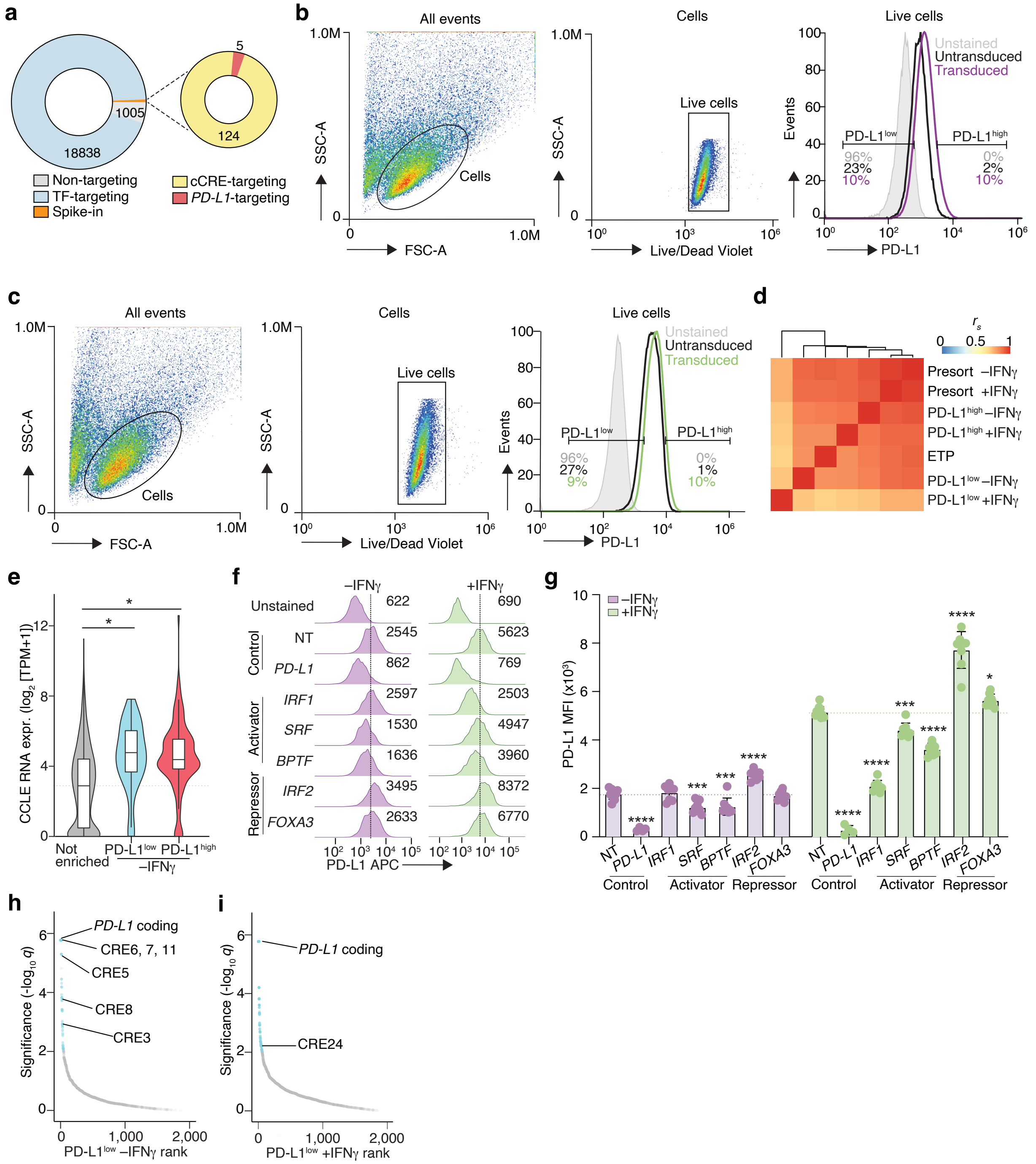

**Supplementary Fig. 4 | TFome-wide CRISPR screen for PD-L1 *trans*-regulators.**
**a,** Guide RNAs in the TFome-wide CRISPR library.
**b, c,** Fluorescence-activated cell sorting gating strategy based on PD-L1 expression for untreated cells (*b*) and IFNγ-treated cells (*c*).
**d,** Spearman correlation of the average normalized gRNA counts in the TFome-wide library-transduced BxPC-3 cell populations ($n$ = 2 biological replicates per cell population).

**e**, Transcript expression of TFs in BxPC-3 cells (Broad Cancer Cell Line Encyclopedia). Significance was determined using a Mann-Whitney *U* test to TFs that were not enriched in either CRISPR screen condition (PD-L1$^{low}$ or PD-L1$^{high}$): *$p < 0.05$.

**f**, PD-L1 surface protein expression after transduction with individual gRNAs targeting *PD-L1*, activator TFs (*IRF1*, *SRF*, and *BPTF*) and repressor TFs (*IRF2* and *FOXA3*) in untreated cells (*left*) and IFNγ-treated cells (*right*). The numbers on the histograms correspond to the mean fluorescence intensity (MFI) of PD-L1. Dashed lines indicate the MFI for non-targeting (NT) control cells.

**g**, MFI of PD-L1 after transduction with individual gRNAs targeting *PD-L1*, activator TFs (*IRF1*, *SRF*, and *BPTF*) and repressor TFs (*IRF2* and *FOXA3*) in untreated cells and IFNγ-treated cells ($n$ = 2 - 4 gRNAs with 2 - 4 biological replicate transductions per gRNA). MFIs are normalized by subtracting the MFI of unstained cells. Significance was determined using a Mann-Whitney *U* test to the non-targeting (NT) group with Dunnett's correction: *$p < 0.05$, **$p < 0.01$, ***$p < 0.001$, ****$p < 0.0001$.

**h, i,** Robust rank aggregation (RRA) of the TFome-wide library with select PD-L1 CRE-targeting perturbations indicated in untreated (*h*) and IFNγ-treated (*i*) PD-L1$^{low}$ flow-sorted cells. FDR-adjusted RRA $q$-values are derived from consistent enrichment of gRNAs targeting the same TF/CRE ($n$ = 10 gRNAs per TF and 4 gRNAs per CRE with 2 biological replicates of each pooled screen).

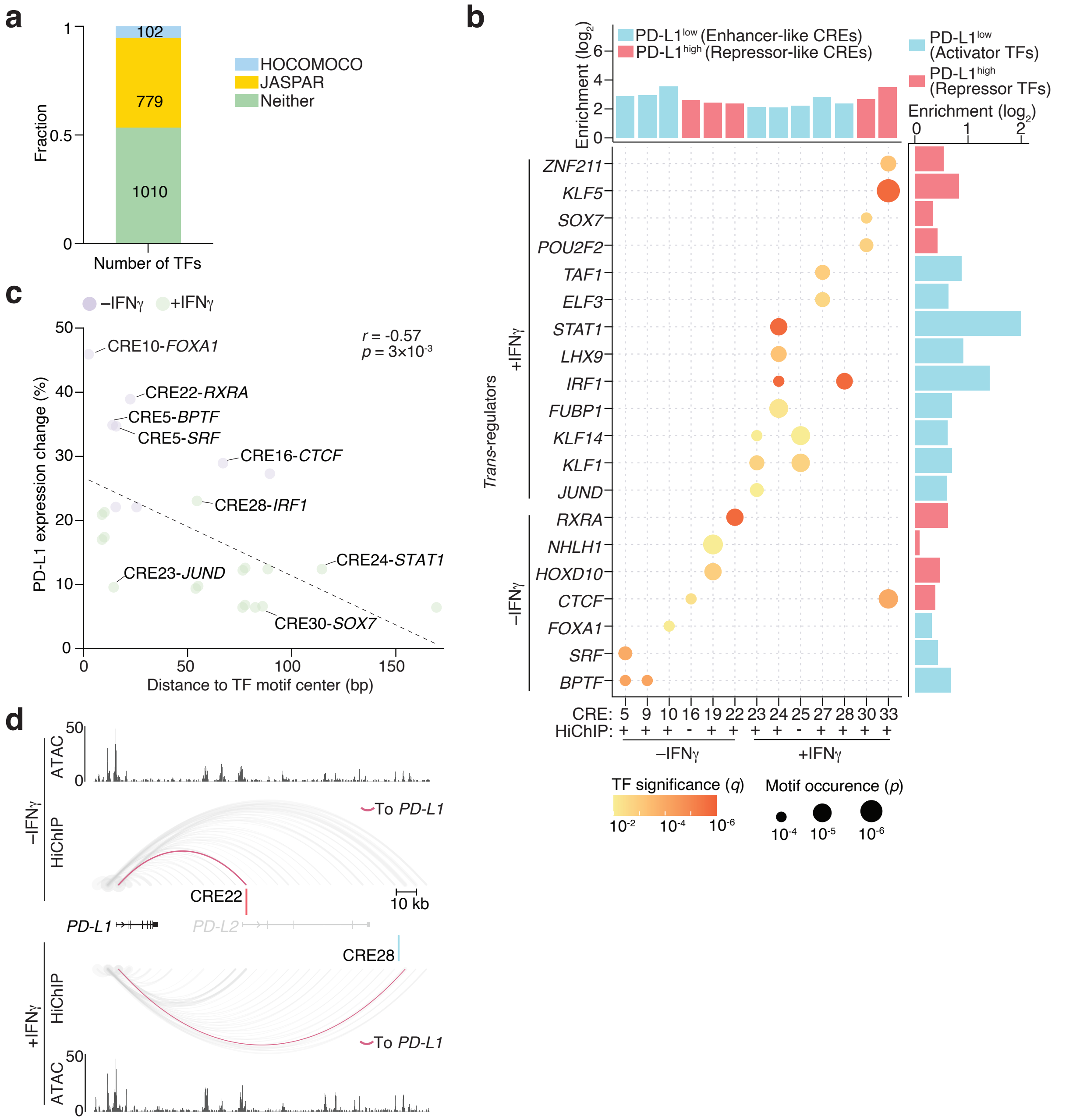


**Supplementary Fig. 5 | Integration of PD-L1 *cis*-regulatory elements and *trans*-regulators.**

**a,** Number of human TFs in the TFome-wide CRISPR screen with TF motifs (position weight matrices) available.

**b**, Motif identification for the 25 predicted CRE-TF regulatory modules. Dot size indicates the significance of motif position-weight matrix matching at each CRE (FIMO log-likelihood *p*). Dot color indicates the significance of the TF in PD-L1 regulation from the corresponding TFome-wide CRISPR screen (FDR-adjusted RRA *q*-values). The upper bar plot shows the enrichment (vs. presort) of CREs from the saturation mutagenesis screen. The bar plot at right shows the enrichment (vs. presort) of TFs from the TFome-wide CRISPR screen. For each CRE, the presence/absence of 3D physical contact (H3K27ac HiChIP loops) to the promoters of *PD-L1* or *JAK2* is denoted at the bottom.

**c,** Pearson correlation between the average change in PD-L1 surface protein expression after CRE perturbation and the average distance between the cut sites of CRE-targeting gRNAs and the center of predicted TF binding site within the CRE ($n$ = 2 gRNAs per CRE and 2 biological replicates per gRNA). This plot includes the same 25 predicted CRE-TF regulatory modules as in panel *b*.
**d,** H3K27ac HiChIP and ATAC-seq signals at repressor-like CRE22 and enhancer-like CRE28. Direct HiChIP loops between CREs and the promoter of *PD-L1* are highlighted.

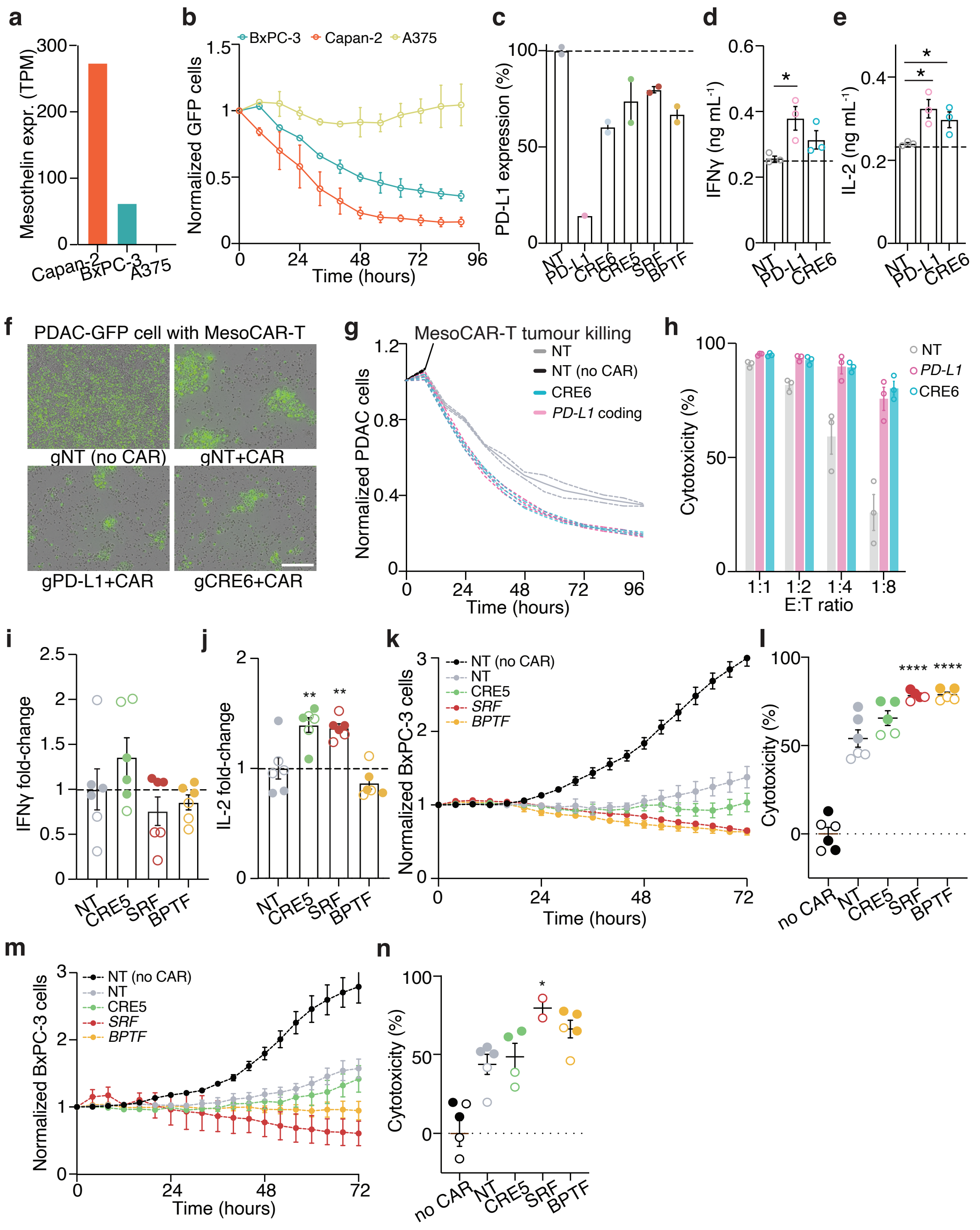


**Supplementary Fig. 6 | PD-L1 *cis*- and *trans*-regulators impact CAR-T mediated immune evasion.**

**a,** Mesothelin transcript expression in PDAC (BxPC-3 and Capan-2) and melanoma (A375).

**b,** BxPC-3, Capan-2 and A375 GFP$^{+}$ cell count when co-cultured with human primary CD8$^{+}$ anti-mesothelin CAR-T (MesoCAR-T) cells at a 1:1 effector-to-target cell ratio ($n$ = 3 biological replicates per cell line).

**c**, PD-L1 cell surface protein expression after transduction with individual non-targeting (NT), *PD-L1* coding exon-targeting, *PD-L1* promoter (CRE6)-targeting and PD-L1 regulator-targeting gRNAs. PD-L1 expression is normalized to NT gRNA-transduced cells ($n = 2$ gRNAs per CRE or TF).

**d, e,** IFNγ (*d*) and IL-2 (*e)* quantification by enzyme-linked immunosorbent assay after 16-hour co-incubation of human primary $CD8^+$ MesoCAR-T cells with Capan-2 PDAC cells transduced with non-targeting (NT), *PD-L1* coding exon-targeting (Coding) or CRE6-targeting gRNAs ($n = 3$ biological replicates per perturbation). Significance was determined using a Mann-Whitney *U* test to non-targeting gRNA-transduced cells with Bonferroni correction: $*p < 0.05$.

**f,** Representative images of BxPC-3-GFP PDAC cells transduced with non-targeting (NT), *PD-L1* coding exon-targeting (coding) or CRE6-targeting gRNAs co-cultured with human primary $CD8^+$ MesoCAR-T cells or untransduced control T cells. Scale bar, 200 µm.

**g,** Counts of BxPC-3-GFP PDAC cells transduced with non-targeting (NT), *PD-L1* coding exon-targeting (Coding) or CRE6-targeting gRNAs when co-cultured with human primary $CD8^+$ MesoCAR-T cells ($n = 3$ biological replicates per perturbation). Cell counts are normalized to the initial timepoint on a per-well basis and all co-cultures are using a 1:1 effector-to-target cell ratio.

**h,** Cytotoxicity of human primary $CD8^+$ MesoCAR-T cells when co-cultured with BxPC-3-GFP cells transduced with non-targeting (NT), *PD-L1* coding exon-targeting (Coding) or CRE6-targeting gRNAs after 96 hours at different effector-to-target cell ratios ($n = 3$ biological replicates per perturbation and cell ratio). Cytotoxicity is normalized to human primary $CD8^+$ cells (no CAR) when co-cultured with BxPC-3-GFP cells transduced with non-targeting (NT).

**i, j,** IFNγ (*i*) and IL-2 (*j)* quantification after 16-hour co-incubation of $CD4^+$ mesoCAR-T cells with PDAC BxPC-3 cells transduced with non-targeting (NT), CRE5-targeting, *SRF*-targeting and *BPTF*-targeting gRNAs ($n = 3$ biological replicates per perturbation and donor). Filled and open circles denote two separate human donors. Significance was determined using a Mann-Whitney *U* test to NT gRNA-transduced cells and Bonferroni-corrected *p*-values are shown: $**p < 0.01$.

**k,** Counts of BxPC-3-GFP cells transduced with non-targeting (NT), CRE5-targeting, *SRF*-targeting or *BPTF*-targeting gRNAs when co-cultured with human primary $CD8^+$ MesoCAR-T cells ($n$ = 2-3 biological replicates per donor for 2 donors). Cell counts are normalized to the initial timepoint on a per-well basis and all co-cultures are using a 1:2 effector-to-target cell ratio.

**l,** Cytotoxicity of human primary $CD8^+$ MesoCAR-T cells when co-cultured with BxPC-3-GFP cells transduced with non-targeting (NT), CRE5-targeting, *SRF*-targeting or *BPTF*-targeting gRNAs after 78 hours at 1:2 effector-to-target cell ratio ($n$ = 2-3 biological replicates per donor for 2 donors). Filled and open circles denote two separate human donors. Cytotoxicity is normalized to human primary $CD8^+$ cells (no CAR) when co-cultured with BxPC-3-GFP cells transduced with a NT gRNA. Significance was determined using a Mann-Whitney *U* test to non-targeting gRNA-transduced cells with Bonferroni correction: $*p < 0.05$, $****p < 10^{-4}$.

**m,** Counts of BxPC-3-GFP cells transduced with non-targeting (NT), CRE5-targeting, *SRF*-targeting or *BPTF*-targeting gRNAs when co-cultured with human primary $CD8^+$ MesoCAR-T cells ($n$ = 2-3 biological replicates per donor for 1-2 donors). Cell counts are normalized to the initial timepoint on a per-well basis and all co-cultures are using a 1:4 effector-to-target cell ratio.

**n,** Cytotoxicity of human primary $CD8^+$ MesoCAR-T cells when co-cultured with BxPC-3-GFP cells transduced with non-targeting (NT), CRE5-targeting, *SRF*-targeting or *BPTF*-targeting gRNAs after 78 hours at 1:4 effector-to-target cell ratio ($n$ = 2-3 biological replicates per donor for 1-2 donors). Filled and open circles denote two separate human donors. Cytotoxicity is normalized to human primary $CD8^+$ cells (no CAR) when co-cultured with BxPC-3-GFP cells transduced with a NT gRNA. Significance was determined using a Mann-Whitney *U* test to NT gRNA-transduced cells with Bonferroni correction: $*p < 0.05$.

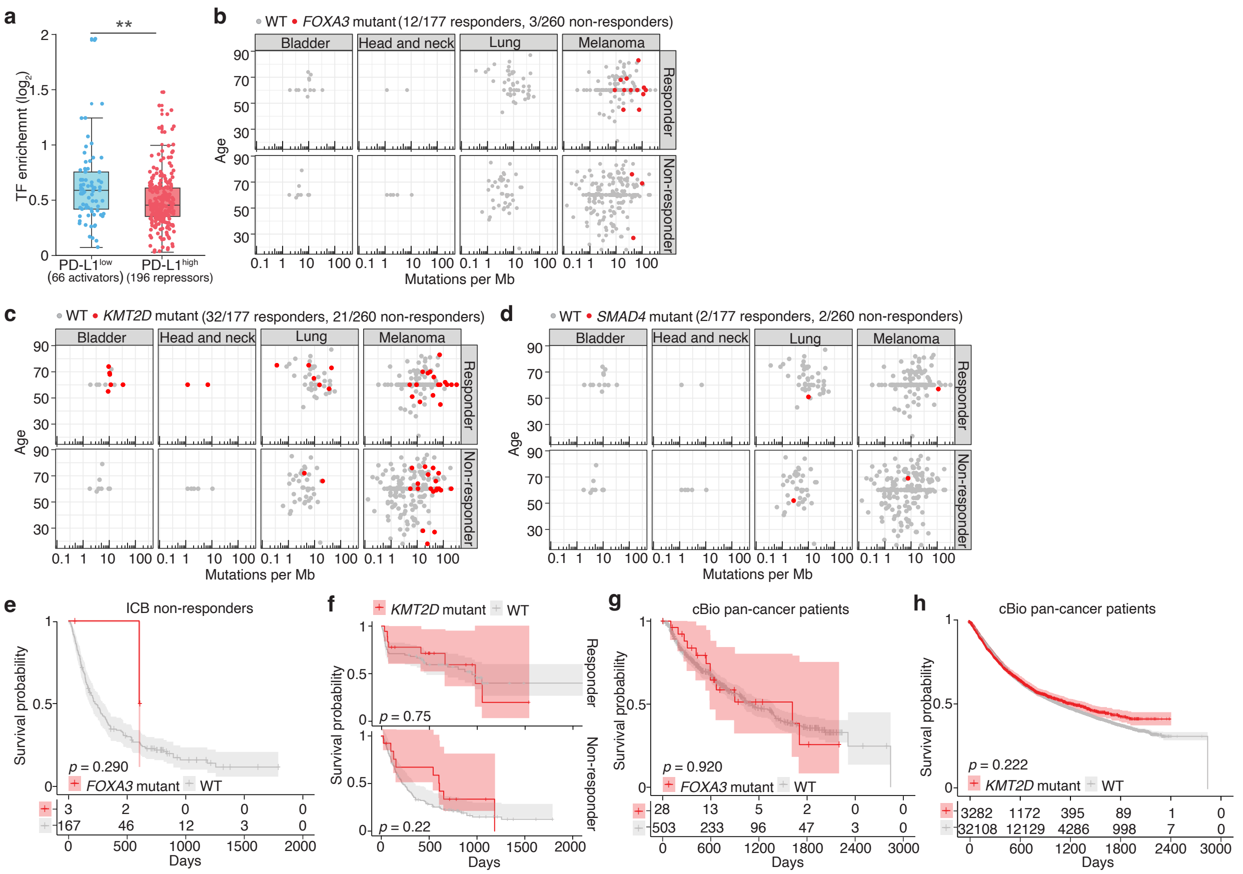


**Supplementary Fig. 7 | Characterization of PD-L1 *trans*-regulators in PDAC patients and immune checkpoint blockade (ICB)-treated patients.**

**a**, Enrichment of top-ranked TFs (RRA $q < 0.01$) in PD-L1$^{low}$ and PD-L1$^{high}$ flow-sorted cells compared to presort cell populations. Significance was determined using a Mann-Whitney *U* test: $^{**}p < 0.01$.

**b-d,** Age, tumor mutational burden (TMB) and tumor type for ICB responders and non-responders with (*red*) and without (*gray*) mutations in *FOXA3* (*b*), *KMT2D* (*c*) and *SMAD4* (*d*) ($n = 437$ patient tumors).

**e,** Overall survival (Kaplan-Meier curve) for ICB non-responders with (*red*) or without (*gray*) mutations in *FOXA3* ($n = 170$ patients with whole-exome sequencing and survival data). Shaded areas indicate the 95% confidence interval and statistical significance was determined using the log-rank test.

**f,** Overall survival (Kaplan-Meier curve) for patients with (*red*) or without (*gray*) mutations in *KMT2D* in ICB responders ($n = 87$ patients with whole-exome sequencing and survival data) (*top*) and ICB non-responders ($n = 170$ patients with whole-exome sequencing and survival data) (*bottom*). Shaded areas indicate the 95% confidence interval and statistical significance was determined using the log-rank test.

**g,** Overall survival (Kaplan-Meier curve) for cBioPortal pan-cancer patients with or without mutations in *FOXA3* ($n = 531$ patients). Shaded areas indicate the 95% confidence interval and statistical significance was determined using the log-rank test.

**h,** Overall survival (Kaplan-Meier curve) for cBioPortal pan-cancer patients with or without mutations in *KMT2D* ($n = 35{,}390$ patients). Shaded areas indicate the 95% confidence interval and statistical significance was determined using the log-rank test.
